# Cap-specific N6-adenosine methyltransferases and applications in m6A Cap-1 mRNA synthesis

**DOI:** 10.64898/2026.10.22.757692

**Authors:** Daniel W. Kneller, Nan Dai, S. Hong Chan

## Abstract

In addition to the 5’ Cap-1 structure, the initiating adenosine of mammalian mRNA can be further modified by PCIF1 (Phosphorylated C-terminal domain-interacting factor 1) to form 2’-O,N6-dimethyladenosine (m6A Cap-1). While the biological role is still under investigation, the implications of m6A Cap-1 in mRNA metabolism and viral-host interactions have spurred interest in PCIF1 and m6A Cap-1 as biomarkers, drug targets, and research tools. Here, we identify and characterize two PCIF1-like cap-specific N6-adenosine methyltransferases, PC04 and PC05 and evaluate their applicability in m6A Cap-1 mRNA synthesis *in vitro.* We found that both enzymes are highly specific to the m7G cap structure and highly active on Cap-1 RNA oligonucleotides and trinucleotide Cap-1 analogs at nanomolar concentrations. We demonstrate that PC04 or PC05 methylation reactions can be incorporated into established post-transcriptional capping and cap analog-based co-transcriptional *in vitro* transcription (IVT) workflows to produce m6A Cap-1 RNAs. Finally, we show that PC05 can be combined with a previously reported FCE::T7 RNA polymerase fusion to synthesize m6A Cap-1 mRNA from DNA templates in a single step, co-transcriptional manner. The robustness of the PCIF1-like cap N6-adenosine methyltransferases can accelerate research into the biological roles of cap m6A modification and development of highly effective mRNA-based therapeutics.

**Graphical Abstract:** 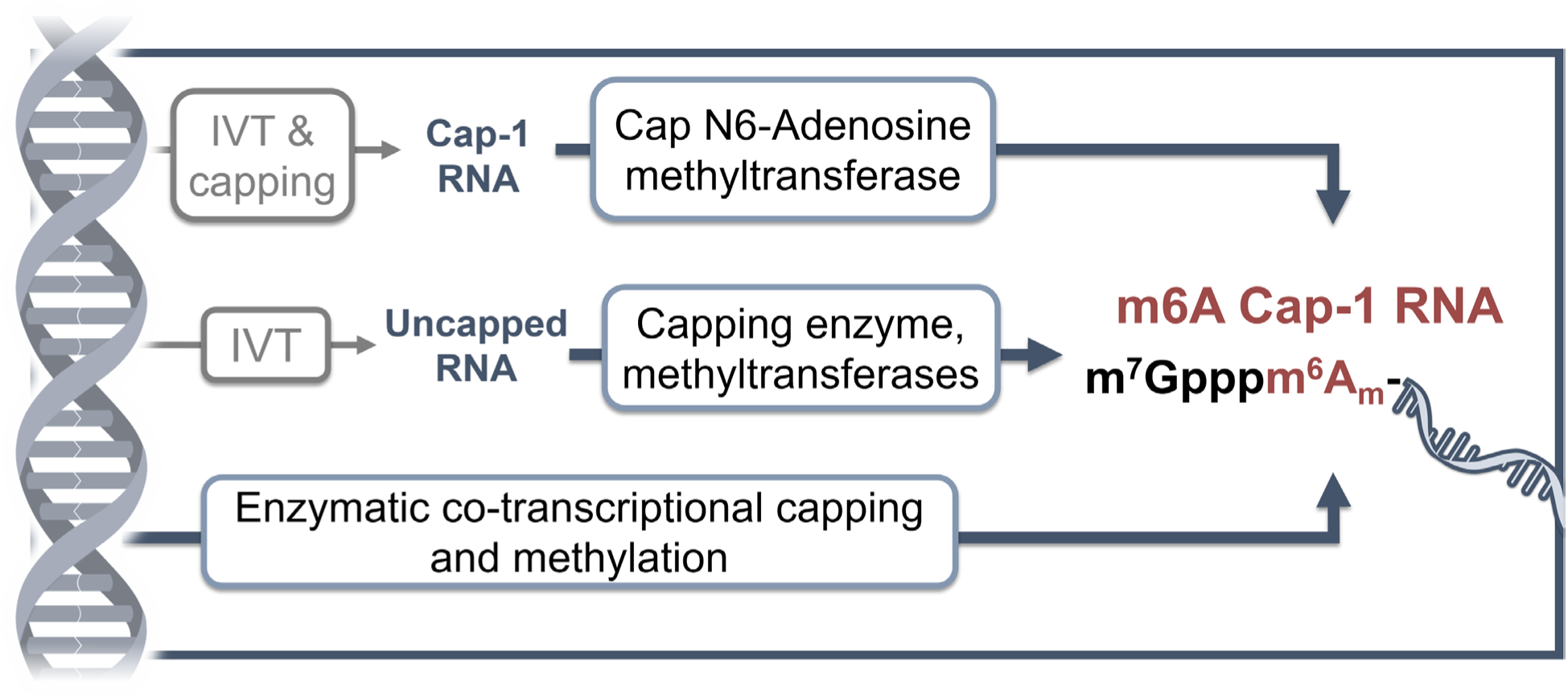

## Introduction

The chemical diversity of mammalian RNA extends beyond the four canonical ribonucleosides through a vast number of chemical modifications. For messenger RNA, modifications can be installed during or after transcription and serve as regulatory elements in localization, metabolism, translation, and innate immune recognition. N6-methyladenosine (m6A) is a prevalent and well-studied modification in human mRNA. The dynamic relationship between m6A writers, erasers, and readers are important for the regulation of essential cellular events and tumorigenesis (1–3). Moreover, chemical modifications of mRNA have been shown to contribute to the safety and effectiveness of mRNA therapeutics and vaccines. The discovery and mechanistic research into the reduced immunogenicity of synthetic mRNA containing modified uridine (pseudouridine and m1-pseudouridine) have contributed to the successful development of mRNA-based vaccines against SARS-CoV-2 infection (4, 5). The potential for nucleotide modifications to achieve favorable properties for mRNA-based therapeutics has driven more efforts in modification detection and research on their effects on cellular biology.

One of the most well-studied mRNA modifications is the 5’ terminal cap of eukaryotic mRNA. The 5’ cap is a critical component in regulating mRNA turnover, nuclear export, translation initiation, and innate immune surveillance. The Cap-0 structure (m7GpppN-) is a N7-methylgunanosine linked by a 5’-5’ triphosphate group to the transcription start nucleotide (TSN). Cap-1 (m7GpppNm-) contains an additional methylation of the TSN ribose at the 2’ oxygen position that is critical for evading innate immune surveillance of the cell. The mammalian Cap-1 structure is installed on nascent mRNA molecules co-transcriptionally via a multi-component enzymatic pathway (FIG 1A).

Multiple cap structures of mammalian mRNA were discovered in human cells in the 1970s (6–8) but their abundance and functions remained unclear until recently. Advancements in mass spectrometry and sequencing methods are now beginning to help reveal the biology and metabolism of the mRNA Cap methylome. For example, the presence of 2’-O-methylation on the second nucleotide of Cap-1 mRNAs (Cap-2 mRNA) is associated with long cytosolic lifetime of specific mRNA molecules (9, 10) suggesting a role of Cap-2 in mRNA turnover (10). The m6A Cap-1, where the Cap-1 structure of adenosine-initiating mRNA is further methylated at the N6-position to form the 2’O,N6-dimethyladenosine structure (m7Gpppm6Am-)(FIG 1B), has been shown recently to be the prevalent form of the 5’ cap of adenosine-initiating mRNA in mammalian cells (11–13).

Despite the prevalence, investigations of the biological role of the m6A Cap-1 structure oftentimes arrive at inconsistent and sometimes contradictory conclusions. It has been shown that the m6A Cap-1 structure confers a mRNA population with increased cellular longevity compared to Cap-1 (13–16) through resistance to DCP2-mediated decapping and reduced susceptibility to microRNA-mediated degradation (14). It was further shown that FTO (Fat mass-associated and obesity-associated protein demethylase), previously known as an internal m6A eraser, demethylates m6A Cap-1 to Cap-1 (14, 17, 18) and reduces the stability of demethylated mRNA molecules (14). However, it has also been reported that the presence of m6Am does not correlate with transcript stability (12, 13, 17, 19). Similarly, the translation level of m6A Cap-1 mRNA relative to Cap-1 transcripts has been reported to increase (12–14) and decrease (19, 20). The translation level of m6A Cap-1 mRNA is thought to be tissue-specific (15). The discrepancies in the literature have been hypothesized to arise from detection methods and gene variant annotation failing to account for the diversity of possible 5′ sequences and cap structures (11). Improvements in detection and profiling methods and systemic studies of the effect of m6A on Cap-1 mRNA transcription and translation will help better understand the biological roles of this 5’ cap modification.

In the cell, producing m6A Cap1 involves a series of catalytic activities (Figure 1A) supplied by enzymes that are co-transcriptionally recruited to the C-terminal domain (CTD) of RNA Polymerase II (Pol II). Originally described as a transcription factor and binder to the CTD of RNA Pol II in 2003 (21), Phosphorylated CTD Interacting Factor 1 (PCIF1) was shown by multiple research groups to be the Cap N6-adenosine methyltransferase that modifies Cap-0 and Cap-1 structures to their respective m6A form (12, 13, 19, 22). The discovery of PCIF1 as the sole cap-specific m6A writer in mammalian cells facilitated investigations into the role of the enzyme and the m6A Cap-1 modification in human diseases. Thus far, PCIF1 and m6A Cap-1 transcripts have been associated with cancer (16, 23–25), viral-host interactions (26–28), parasites (29), and obesity (30). To date, m6A Cap-1 has only been reported in vertebrate and viral RNA but homologs of PCIF1 have been reported in insects (15, 31) and trypanosomes (15).

**Figure 1.**
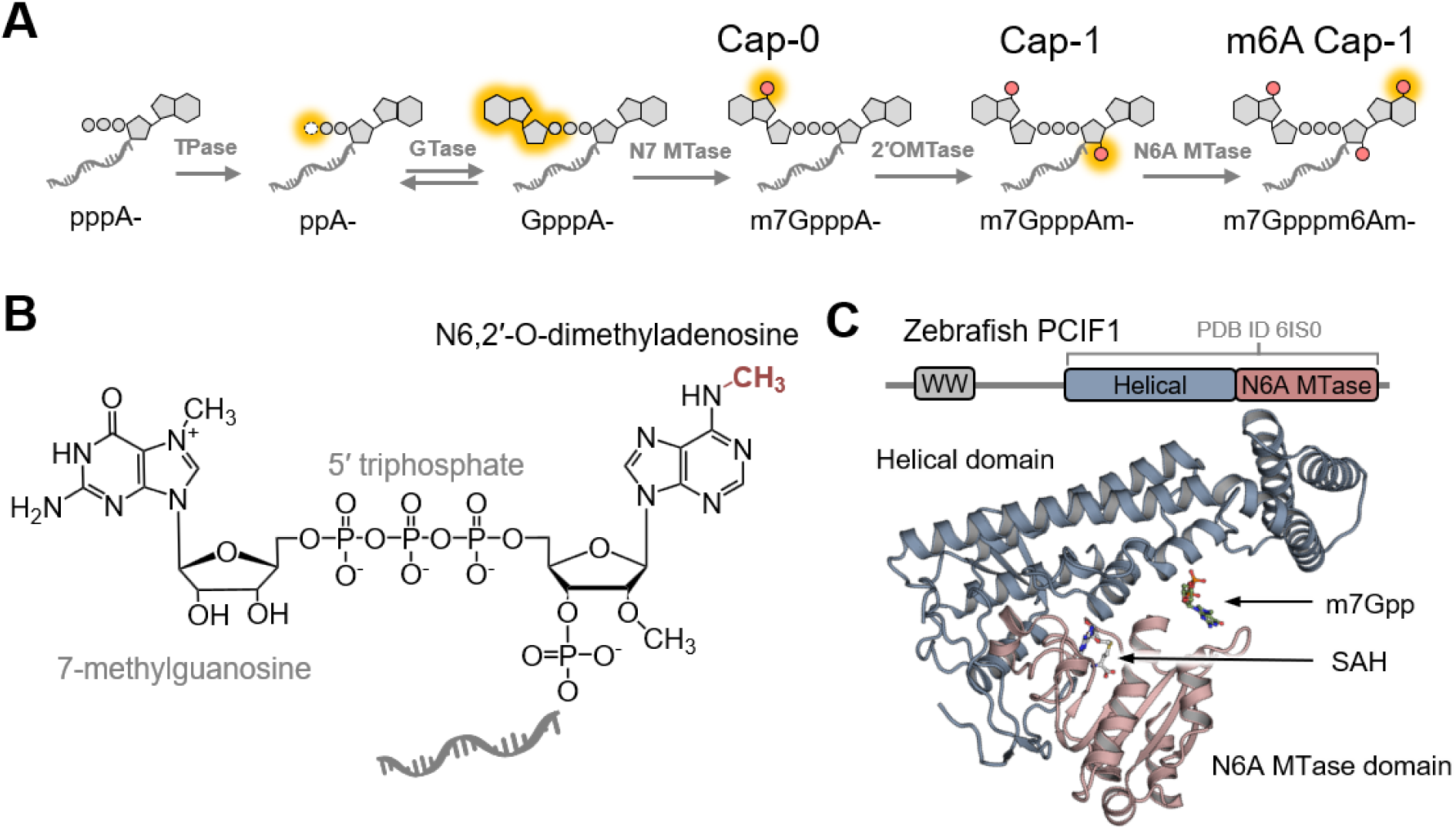
Biosynthetic pathway of m6A Cap-1 installation to message RNA. **(A)** Enzymatic pathway to install the m6A Cap-1 structure to the 5’ end of RNA found in nature. The triphosphate group at the 5’ end of a nascent transcript is hydrolyzed to form a diphosphate group by RNA triphosphatase (TPase). The reversible RNA guanylyltransferase (GTase) adds a GMP to the diphosphate group to form a guanosine 5’-5’ triphosphate group at the 5’ end of the RNA. The cap N7-guanosine methyltransferase (N7 MTase) then adds a methyl group to the N7 position of the guanosine cap to form the Cap-0 structure. A cap 2’-O-methyltransfease (2’OMTase) then adds a methyl group to the 2’-O position of the ribose of the first nucleotide to form the Cap-1 structure. Finally, a cap-specific N6-adenosine methyltransferase adds a methyl group to the initiating adenosine at its N6 position, forming the m6A Cap-1 structure. The addition and subtraction of chemical groups in each reaction are highlighted in yellow. The methyl groups are highlighted in red. **(B)** The chemical structure of the m6A Cap-1 structure. The m6A methyl group is highlighted in red. **(C)** The Zebrafish PCIF1 domain organization and atomic structure with bound SAH and a partial m7G cap (PDB ID 6IS0).

**Figure 2.**
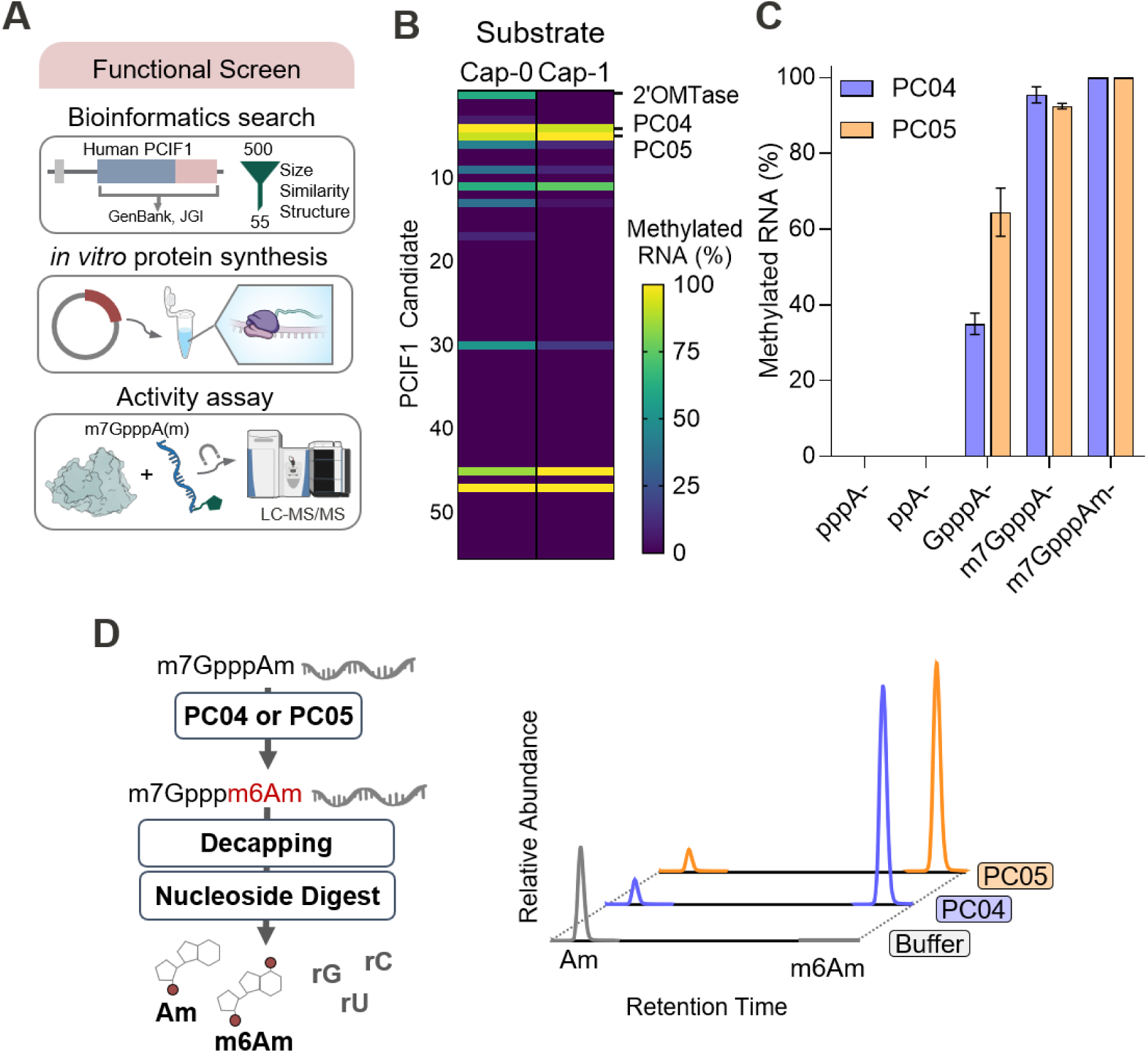
Discovery and substrate specificity of non-metazoan cap-specific N6-adenosine methyltransferase. **(A)** The workflow of the functional screening. A bioinformatic search of the JGI Integrated Microbial Genomes & Microbiomes was carried out for proteins homologous to human PCIF1 by applying filters on the size, sequence similarity and structural elements. The selected candidates were synthesized using an *in vitro* protein synthesis kit (PURExpress ΔRF123, New England Biolabs) and directly tested in an RNA methylation assay. The resulting RNA was then enriched and analyzed using intact LC-MS/MS. **(B)** A representative heat map of the functional screening results. The level of methylation is colored with the Viridis color scale. Each row represents the results of a selected candidate protein. The assay was performed on RNA oligonucleotides containing a Cap-0 or Cap-1 structure. Five candidates, including PC04 and PC05, exhibited high levels of methylation activity on both substrates. As expected, Vaccinia cap 2’OMTase used as a positive control is active only Cap-0 RNA substrate. **(C)** Substrate specificity of the methylation activity of PC04 and PC05. Both enzymes exhibit high levels of activity on Cap-0 (m7GpppA-) and Cap-1 (m7GpppAm-) RNA but lower activity on G-capped (GpppA-) RNA. No methylation was detected on 5’ triphosphate or diphosphate RNA. **(D)** Verification of the chemical structure of the methylation product. A synthetic 25mer with the Cap-1 structure (m7GpppAm-) contains a single cap-adjacent 2’-O-methyl-adenosine (Oligo 14) was used as substrate. After decapping and digestion into nucleosides, UHPLC-MS/MS analysis with dynamic multiple reaction monitoring (dMRM) indicated that m6Am was only detected when PC04 or PC05 was present. Details of the elution times and the results of dMRM can be found in Supplementary FIG S3.

**Figure 3.**
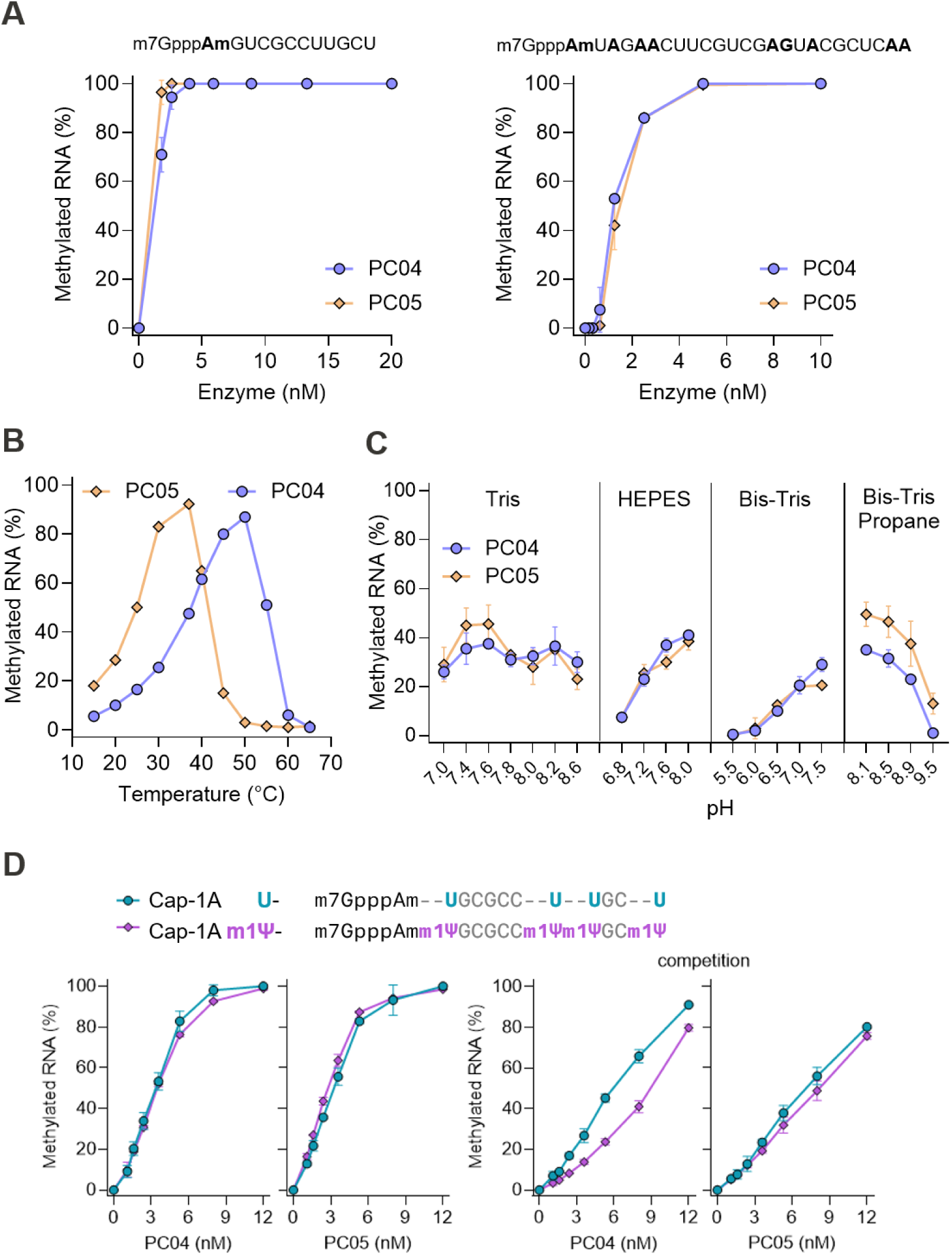
Biochemical characterization of PC04 and PC05. **(A)** PC04 and PC05 are highly active at nanomolar concentrations. Decreasing concentrations of PC04 and PC05 were incubated with 1 µM of indicated Cap-1 RNA oligos. LC-MS/MS analysis shows that 5 nM of both enzymes methylate 100% of a 12-nt Cap-1 RNA oligonucleotide (Oligo 1) containing a single cap-adjacent adenosine (left panel) and a 25-nt RNA oligonucleotide containing an additional 7 internal adenosines (Oligo 6, right panel) after 30 min incubations at 37°C. Consistent with the substrate specificity results above, only one methylation event was detected on the 25-nt Cap-1 RNA containing multiple internal adenosines. **(B)** Temperature range. One micromolar of a Cap-1 RNA oligonucleotide containing a single adenosine at +1 position (Oligo 1) was incubated with 2 nM of PC04 or PC05 at temperatures 15-65°C for 30 min. LC-MS/MS analysis showed that PC04 methylated 50% or more of the Cap-1 substrate from 40 to 60°C (blue trace). PC05 methylated 60% or more of the Cap-1 substrate from 30 to 40°C (orange trace). **(C)** pH range. The methylation activity of PC04 and PC05 on Oligo 1 were assessed in buffers consisting of Tris, HEPES, Bis-Tris, or Bis-Tris Propane (50 mM) at appropriate pH values. LC-MS/MS analysis shows that both enzymes exhibited highest methylation activity at pH 7.0-8.0 in Tris and HEPES buffers and reduced activity at pH values below ∼6.5. PC05 retained noticeable activity up to pH 9.5 at which PC04 is inactive. **(D)** Methylation activity on m1Ψ RNA. PC04 (left panel) or PC05 (right panel) was incubated with 1 µM of a Cap-1 12mer RNA (Oligo 12) containing a single 2’-O-methyl-adenosine at the +1 position and either four unmodified uridines (Cap-1 AU; Oligo 12) or N1-methylpseudouridines (Cap-1 Am1Ψ 12mer; Oligo 13) at 37°C for 30 min. LC-MS/MS analysis shows that PC04 methylates both RNAs equally but prefers unmodified RNA when both RNAs are present (competition). PC05, on the other hand, methylates both RNAs equally separately or simultaneously. Datapoints are average values from duplicated experiments. Error bars represent one standard deviation.

**Figure 4.**
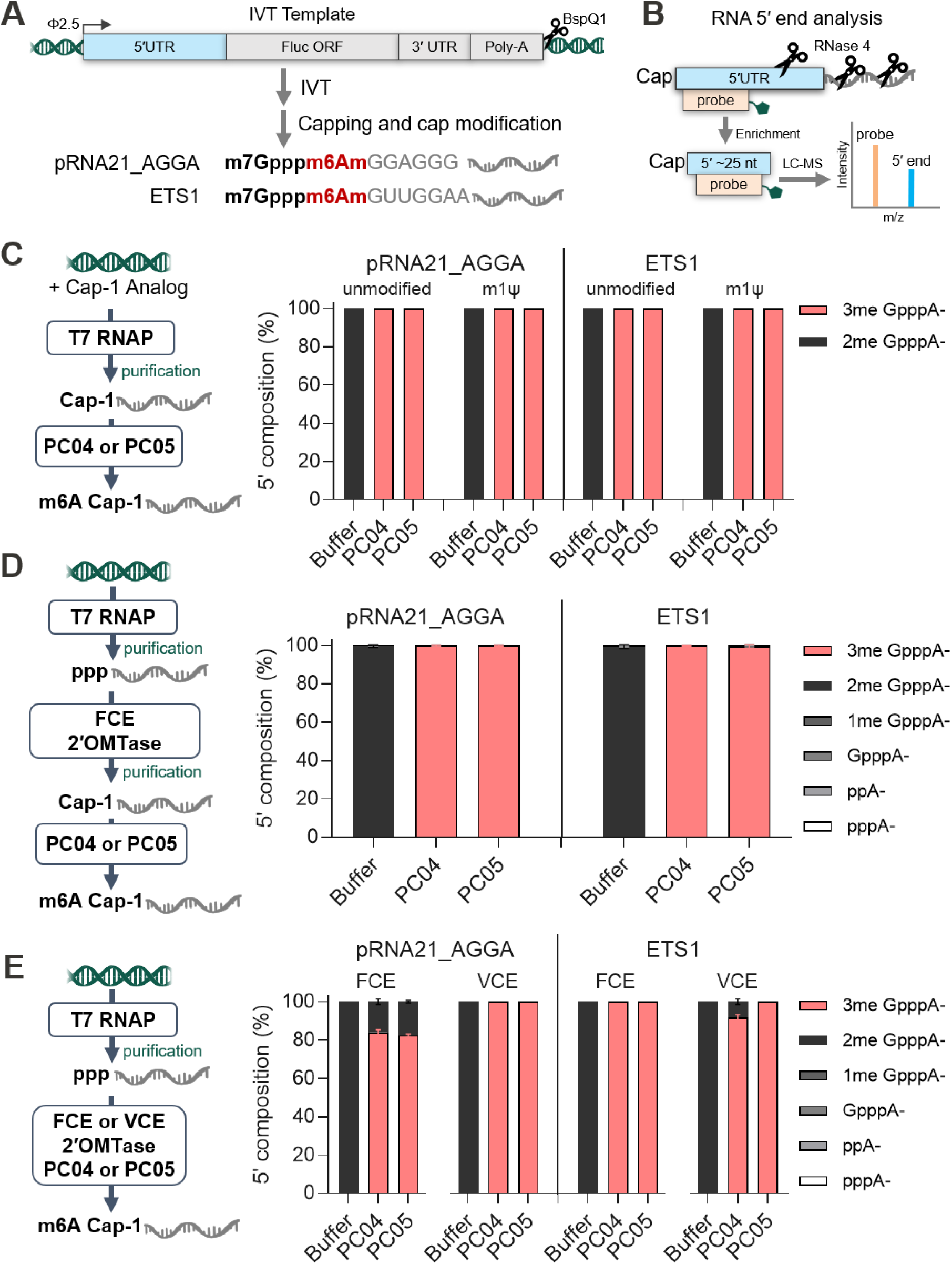
Application of PC04 and PC05 in post-transcriptional workflows. **(A)** A schematic of experimental design and setup. The major components of the IVT templates (from 5’ to 3’ direction) are T7 Phi2.5 promoter, a 5’ UTR (pRNA21_AGGA or ETS1), a firefly luciferase ORF, a 3’ UTR, a poly(A) tail and a BspQI restriction site. **(B)** RNA 5’ end analysis. After annealing to a DNA probe complementary to the first 18-25 nt, the IVT RNA was subjected to RNase 4 cleavage. The double-stranded region resistant to RNase 4 was enriched and analyzed by intact LC-MS/MS. **(C)** PC04 and PC05 modify IVT RNA co-transcriptionally capped using trinucleotide cap analogs. Using trinucleotide cap analog CleanCap^®^ AG, the co-transcriptionally capped IVT RNAs containing uridine or m1Ψ were purified and subjected to PC04 or PC05 reactions containing 120 nM of enzyme. LC-MS/MS analysis indicated that 100% of the modified or unmodified RNAs were methylated by PC04 and PC05 and carried the m6A Cap-1 structure (3me GpppA-) RNAs in the control reactions (buffer) carried the Cap-1 structure (2me GpppA). **(D)** PC04 and PC05 modifies IVT RNA enzymatically capped by FCE and modified by Vaccinia 2’OMTase. After IVT, the RNAs were purified and subjected to Cap-1 incorporation using FCE and Vaccinia 2’OMTase. After purification, the Cap-1 RNAs were subjected to PC04 and PC05 reactions containing 120 nM of enzyme. LC-MS/MS indicated that 100% of the PC04-or PC05-treated RNAs carries the m6A Cap-1 structure (3me GpppA-). RNAs in the control reactions (buffer) carried the Cap-1 structure (2me GpppA). **(E)** PC04 and PC05 in one-pot reaction containing FCE/VCE and Vaccinia 2’OMTase. IVT RNAs were purified and subjected to one-pot RNA capping and modification reactions containing 1 U/µL of FCE or VCE, 5 U/µL vaccinia 2’OMTase and 120 nM of PC04 or PC05. LC-MS/MS analysis indicated that PC04 and PC05 modified 80% of Cap-1 pRNA21_AGGA RNA capped by FCE but 100% of the same RNA modified by VCE. Both PC04 and PC05 modified 100% of Cap-1 ETS RNA capped by FCE but PC04 modified only ∼90% of the same RNA when capped by VCE while PC05 modified 100% of the RNA under the same conditions.

The metazoan PCIF1 tertiary structure possesses three domains (12). X-ray crystal structures of the human and zebrafish PCIF1 constructs (FIG 1C) confirm a class I methyltransferase domain with a conserved Rossman Fold for binding the methyl-doner *S*-adenosyl methionine (SAM) and a canonical catalytic motif, [DN]-P-P-[YFW], conserved in DNA and RNA N6-adenosine methyltransferases (12, 32). Cap specificity appears to be conferred by a distinct helical domain that orients the substrate m7G cap and TSN adenosine nucleobase into their respective binding pockets within the methyltransferase domain (12). Human PCIF1 possesses a positively charged groove that is postulated to function as a binding surface for RNAs with a minimum six nucleotides in length (12). The WW domain near the N-terminus interacts with the RNA Pol II CTD, which also recruits the RNA capping and cap modifying enzymes responsible for Cap-1 formation in a phosphorylation state dependent manner (33, 34), facilitating an ordered co-transcriptional formation of m6A Cap-1 (12).

The Cap-1 structure is critical for pharmacological efficacy of synthetic mRNAs by reducing immunogenicity, increasing stability, and facilitating efficient protein translation. Incorporation of modifications such as N6-methylation on adenosine-initiating mRNAs is one of the strategies to improve efficacy of synthetic mRNA. Synthetic mRNAs are generally synthesized by *in vitro* transcription (IVT) where a DNA template containing an appropriate promoter sequence and the target mRNA sequence is transcribed by the corresponding DNA-dependent RNA polymerase. An appropriate 5’ cap structure can be incorporated into the IVT transcript by established methods such as post-transcriptional capping using mRNA capping enzymes from Vaccinia virus (VCE) or Faustovirus (FCE) (35) in combination with Vaccinia cap 2’-O-methyltransferase or by co-transcriptional capping using a trinucleotide Cap-1 analog in IVT.

The use of chemically synthesized m6A trinucleotide Cap-1 analogs co-transcriptionally during IVT is a commonly used method to synthesize m6A Cap-1 RNA (36). This method, however, requires modification of IVT conditions such as high concentrations of m6A Cap-1 analogs and T7 RNA polymerase (36, 37). The demanding conditions represent a barrier to the research on the biological roles and therapeutic application of the m6A Cap-1 structure. Enzymatic methods, on the other hand, are usually less resource demanding and may facilitate more research on the cap structure. We therefore set out to discover and characterize a robust cap-specific N6A methyltransferase and evaluate its applicability in m6A Cap-1 mRNA synthesis.

In this study, we identified and characterized the biochemical properties of two PCIF1 homologs that are specific to the 5’ cap structure of adenosine-initiating RNA. We further establish methods that can be used to incorporate the PCIF1 homologs into existing IVT workflows to synthesize m6A Cap-1 mRNA *in vitro* post-transcriptionally and co-transcriptionally. We further showed that the PCIF1 homologs can modify trinucleotide Cap-1 analogs to generate m6A Cap-1 analogs commonly used in co-transcriptional capping workflows. The application of a robust RNA cap N6A methyltransferase in m6A Cap-1 RNA synthesis can help accelerate the research on the biological role of m6A Cap-1 and the development of more effective mRNA therapeutics.

## Results

### Functional screening identifies Cap m6A methyltransferases from non-metazoan organisms

To identify a N6-adenosine methyltransferase with 5’ capped TSN specificity, protein sequences were selected from a bioinformatics search and then functionally screened for enzyme activity, shown schematically in FIG 2A. A list of 500 non-redundant PCIF1-like ORFs were aggregated from BLAST searches of the human PCIF1 combined methyltransferase-helical domain module as a query against non-metazoan organisms and metagenome databases. The collection was filtered by excluding ORFs >480 amino acids and those with >85% similarity based on BLOSUM62 scoring. The tertiary structure of the retained ORFs were computationally predicted (38, 39) and compared to the zebrafish enzyme-substrate complex X-ray crystal structure (12). ORFs without the N6-adenosine methyltransferase catalytic motif and a cap-stacking tryptophan residue in the structural predictions were discarded. A final curated list of 55 putative PCIF1 ORFs were selected for functional assays.

Each of the PCIF1-like N6-adenosine methyltransferase candidates was expressed *in vitro* and evaluated by an *in vitro* functional assay. To achieve high throughput and low nuclease contamination, protein expression was done using a reconstituted *E. coli* transcription-translation system *in vitro* (40) and the translation products were directly evaluated for methyltransferase activity. Twenty-five nucleotide long single stranded RNA oligos (25mers) containing a 5′ Cap-0 or Cap-1 structure, a single adenosine at the TSN, and a 3′-desthiobiotin enrichment tag were used as substrates (Supplementary Table S1). The level of methylation was assessed using liquid chromatography–mass spectrometry (LC-MS). A plasmid encoding *Vaccinia* virus cap 2′-O methyltransferase served as a control for *in vitro* expression and cap-specific RNA methyltransferase activity. Out of the 55 candidates, 10 displayed methyltransferase activity on at least one of the two capped RNA oligos. Five candidates exhibited a majority abundance of the methylated products on both substrates (FIG 2B). However, only <u>P</u>CIF1 <u>C</u>andidate 04 and 05 (PC04 and PC05) were amenable to soluble bacterial expression and purification from *E. coli* (Supplementary FIG S1).

Substrate specificity of PC04 and PC05 was evaluated using a panel of RNA oligos containing internal adenosines or a single adenosine as TSN with 5′ groups normally present in *in vitro* transcripts before and after undergoing enzymatic capping (Supplementary Table S1). Both enzymes are highly specific for the 5’ cap structure – while uncapped transcripts (pppA- or ppA-) were not methylated, the Cap-1 RNA was completely methylated; the Cap-0 RNA was methylated to ∼90%; the unmethylated Cap (GpppA-) was methylated to 35% and 65% by 10 nM of PC04 and P05, respectively (FIG 2C). Furthermore, no methylation was observed on RNA oligos sporting hydroxyl 5′ ends (HO-) with 2’-O-methyladenosines located terminally or internally (Supplementary FIG S2). To verify the position of methylation, PC04 and PC05 reaction products on a Cap-1 RNA oligo containing a single adenosine as the TSN were decapped using a mRNA decapping enzyme and then subjected to nucleoside analysis by LC-MS/MS with dynamic multiple reaction monitoring (dMRM). Significant amounts of N6,2’-O-dimethyladenosine (m6Am) on the TSN were detected only in the presence of PC04 or PC05 (FIG 2D, Supplementary FIG S3). Details of the elution times and the results of dMRM can be found in Supplementary Information.

### Biochemical characterization of PCIF1-like enzymes on RNA oligos

The biochemical properties of PC04 and PC05 were investigated by *in vitro* assays. An endpoint enzyme activity assay was first used to establish an enzyme concentration range for subsequent assays. The level of Cap-1 RNA oligo substrate and methylated product in the quenched enzymatic reactions were measured by LC-MS analysis. Micromolar concentrations of two different Cap-1 RNA substrates (containing a single adenosine as the cap-adjacent nucleotide or multiple internal adenosines) were 100% methylated after 30 min incubations at 37°C with low nanomolar concentrations of PC04 or PC05 (FIG 3A). Consistent with the substrate specificity results above, only one methylation event was detected on the Cap-1 RNA containing multiple internal adenosines.

We determined the temperature range for PC04 and PC05 methylation activity by incubating reactions with Cap-1 RNA oligos at 15-65°C at limiting enzyme concentration (2 nM). PC04 methylated >50% of the RNA oligos at temperatures between 40-55°C with noticeable activity observed as low as 25°C and as high as 60°C (FIG 3B). PC05 methylated >50% of the RNA oligos at 25-40°C with noticeable activity below 20°C and greater than 45°C. Hence, the optimal temperature for methylation activity under the conditions tested was 50°C and 37°C for PC04 and PC05 respectively.

The effective pH range for PC04 and PC05 was determined by assessing methylation activity of the enzymes on Cap-1 RNA oligos in 50 mM Tris, HEPES, Bis-Tris, or Bis-Tris Propane buffers adjusted to the appropriate pH values (FIG 3C). In general, PC04 and PC05 exhibited the highest methylation activity at pH 7.0-8.0 in Tris or HEPES buffers and reduced product accumulation at pH values below neutral. PC05 displayed residual activity in alkaline Bis-Tris Propane conditions up to pH 9.5 where PC04 was inactive. To gauge the performance of the two enzymes under IVT pH regime, which is commonly maintained between pH 7-8 using a Tris buffer, the methylation activity of PC04 and PC05 at 3 different enzyme concentrations was tested incrementally in Tris buffer at pH 7.0 to 8.4. Both enzymes displayed steady methylation activity across this range (Supplementary FIG S4).

Given the importance of the N1-methylpseudouridine (m1Ψ) modification in therapeutic applications, the effect of m1Ψ-modifications on PC04 and PC05 activity was examined by comparing their activity on RNA substrates with or without m1Ψ substitution (FIG 3D). Synthetic Cap-1 12mer RNA oligonucleotides with the canonical uridines or m1Ψ at the +2 position and 3 other distal positions were incubated with a PC04 or PC05 dilution series. Under these conditions, 12 nM of PC04 and PC05 efficiently methylated more than 98% of both modified and unmodified RNA. To help understand if the enzymes exhibit selectivity between uridine- and m1Ψ-containing RNA, a competition experiment was performed by supplying PC04 or PC05 with an equimolar mixture of the unmodified and modified 12mers. Here, PC04 converted more unmodified substrate than m1Ψ-modified RNA at low nanomolar enzyme concentrations. In contrast, PC05 methylated both RNAs to the same extent over the enzyme concentration range tested (FIG 3D).

Biochemical characterization demonstrates that PC04 and PC05 exhibit highly specific and robust catalytic activity on m7G-capped RNA oligos. While PC04 and PC05 exhibit maximum methylation activity between pH 7-8 and at 50°C and 37°C, respectively, both enzymes are highly active at temperatures regularly used in biochemical processes (25-37°C). Finally, both enzymes efficiently methylated Cap-1 oligos with m1Ψ substitutions adjacent to the TSN. Competition experiments suggest that PC04 may prefer RNA containing uridine over m1Ψ, and that PC05 displayed no observable discrimination.

### PCIF1-like enzymes methylate Cap-1 RNA and integrate with one-pot capping with high efficiency

To verify methylation activity of PC04 and PC05 on Cap-1 RNA synthesized by IVT, we first assessed their utility as simple appendments to conventional capped mRNA IVT workflows. Plasmid DNA were constructed and linearized to serve as IVT templates for model mRNA transcripts with two different 5′ UTR sequences followed by a Firefly luciferase (FLuc) coding sequence and a 3′UTR-60nt Poly-A tail (∼2.1 Kb) (FIG 4A). A 5′ sequence derived from pRNA21 with a 5′ extension of AGGA (pRNA21_AGGA) served as a 5′ UTR unrelated to known PCIF1 target substrates. The 5′ UTR of human enhancing transcription factor proto-oncogene 1 transcription factor (ETS1) was selected as a model for a verified endogenous human PCIF1 substrate sequence (27). Composition of capping and cap methylation reaction products were analyzed using a nuclease protection-based method (41) (FIG 4B). In this assay the 5′ sequence of the target RNA is hybridized to a biotinylated DNA oligonucleotide to prevent endonuclease digestion by the single-strand site-specific human RNase 4. The protected 5′ end fragments are enriched then analyzed by intact LC-MS/MS. The mass of RNase 4 cleavage fragments allows the differentiation of the relevant 5’ cap structures (pppA-, ppA-, GpppA-, and GpppA-with 1, 2 or 3 methyl groups) where 1Me GpppA is annotated as Cap-0, 2Me GpppA as Cap-1 and 3Me GpppA-as m6A Cap-1.

PC04 and PC05 were first tested on purified pRNA21_AGGA and ETS1 Cap-1 RNA prepared using IVT with the trinucleotide Cap-1 analog m7GpppAmG (FIG 4C). Cap-1 RNA transcripts with and without m1Ψ-modification were 100% methylated after incubation with 120 nM of either enzyme while no methylation was detected in the absence of enzyme (Buffer). The same result was achieved for purified Cap-1 mRNA prepared using post-transcriptional enzymatic capping with Faustovirus Capping Enzyme (FCE) and Vaccinia 2′-O methyltransferase (2′OMTase) (35) (FIG 4D). These results demonstrate that PC04 and PC05 can be used to install the Cap m6A modification on Cap-1 RNA synthesized using established methods.

To further streamline enzymatic incorporation of m6A Cap-1 to synthetic mRNA, purified IVT transcripts (containing unmodified uridine and a 5′-triphosphate group, pppA-) were incubated with 2′OMTase and capping enzymes from *Faustovirus* (FCE) or *Vaccinia* (VCE) in a one-pot format. The presence of 120 nM PC04 or PC05 in either one-pot reactions resulted in RNAs with a majority 5′ composition consistent with m6A Cap-1 (FIG 4E). PC04 or PC05 in the presence of FCE and 2′OMTase generated ∼80% capped pRNA21_AGGA transcripts with the m6A Cap-1 modification, whereas in the one-pot VCE-based reactions both enzymes achieved 100% m6A Cap-1 RNA. For the ETS1 transcript, PC04 and PC05 achieved 100% methylation efficiency in conjunction with FCE and 2′OMTase. In the presence of VCE and 2′OMTase, PC04 and PC05 fully modified >90% and 100% of the EST1 transcript, respectively. Although exhibiting variability on different RNA sequences and capping enzyme partners, PC04 and PC05 can be readily incorporated into one-pot post-transcriptional enzymatic capping methodologies to install the m6A Cap-1 structure to synthetic RNA molecules.

### PCIF1-like enzymes efficiently methylate trinucleotide Cap-1 analogs

The human PCIF1 enzyme was reported to require a minimum length of 6 nucleotides for efficient methylation of the RNA molecules (12). Assuming a similar property, trinucleotide Cap-1 analogs would not be substrates of the PCIF1-like enzymes and the enzymes would methylate the newly synthesized Cap-1 RNA generated by co-transcriptional capping using trinucleotide cap analogs in a one-pot setting. To our surprise, 200 nM of PC04 or PC05 only resulted in <10% N6A methylation in the RNA synthesized (FIG 5A). These reactions are formulated to maximize capped RNA yield using high NTP and cap analog concentrations. To investigate if such conditions inhibit PC04 or PC05 methylation activity, we performed competition assays where PC04 and PC05 were incubated with a Cap-1 RNA oligo in the presence of ribonucleoside triphosphates (NTPs) up to 10 mM (FIG 5B). PC04 RNA methylation activity was reduced at NTP concentrations of 2.5 mM and higher. However, PC05 activity was unaffected by up to 10 mM of individual NTPs, suggesting that high NTP concentrations in IVT conditions is not a factor in low methylation activity of PC05.

We next investigated the effect of trinucleotide Cap-1 analogs on PC04 and PC05 activity. In a similar competition assay, increasing concentrations of trinucleotide Cap-1 analog AG (PubChem CID: 164886585) or AU (PubChem CID: 155923781) were added to methylation reactions of a Cap-1 RNA oligonucleotide containing 10 nM PC04 or PC05. We observed a reduction of methylated RNA products dependent on trinucleotide Cap-1 analog concentration for both enzymes (FIG 5C). The fact that PC05 that is not inhibited by up to 10 mM of NTP exhibits a similar level of methylation as PC04 (inhibited by both NTPs and Cap-1 analogs AG and AU) suggests that trinucleotide Cap-1 analogs exhibit a stronger inhibitory effect than NTPs on the enzymes.

To further investigate if trinucleotide Cap-1 analogs were substrates for PC04 and PC05, Cap-1 analog AG (1 µM) was reacted with varied concentrations of PC04 or PC05. Intact LC-MS/MS analysis shows that the Cap-1 analog is methylated in an enzyme concentration dependent manner (FIG 5D). PC05 exhibits a higher level of methylation than PC04 at low nanomolar concentrations.

Millimolar concentrations of trinucleotide Cap-1 analogs are commonly used in co-transcriptional capping IVT. With the robust methylation activity of PC04 and PC05 on micromolar concentrations of trinucleotide Cap-1 analogs, we further investigated whether the enzymes can be used to convert trinucleotide Cap-1 analogs to their m6A version in millimolar scale. Reactions containing 4 or 8 mM of trinucleotide Cap-1 analog AG were incubated with PC04 or PC05 overnight and analyzed for their methylation status (Fig 5E). Under these conditions, PC04 methylated >99% and ∼60% of 4 mM and 8 mM Cap-1 analog AG, respectively while PC05 methylated >99% the Cap-1 analog at both substrate concentrations (FIG 5E).

**Figure 5.**
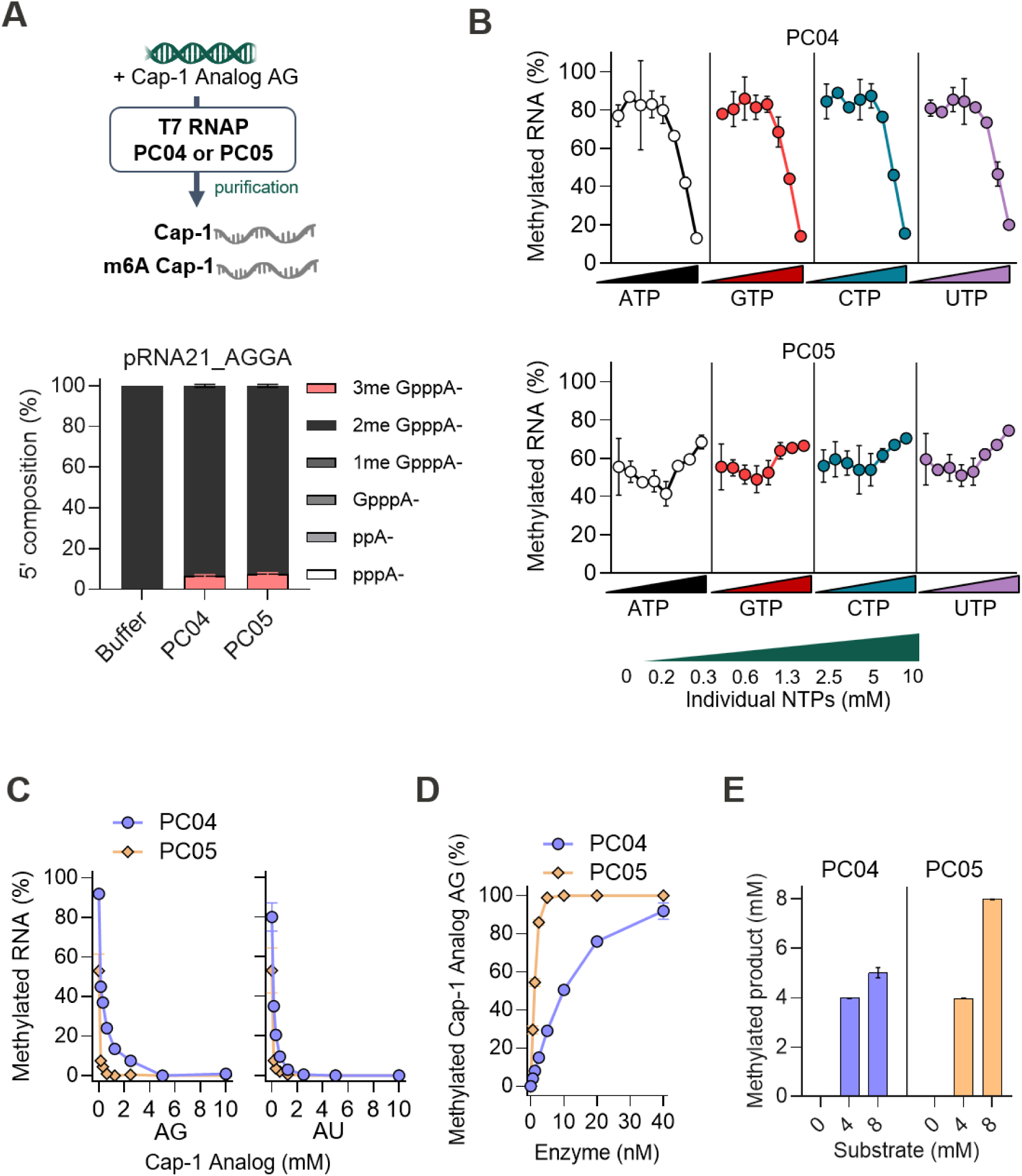
Methylation activity on trinucleotide cap analogs. **(A)** One-pot trinucleotide cap analog co-transcriptional capping and N6-adenosine methylation reactions. A schematic of the workflow (top panel) and RNase 4-LC-MS/MS 5’ cap analysis (bottom panel). PC04 and PC05 only achieved very low levels of methylation (>10%) under these conditions. **(B)** High NTP concentrations inhibit PC04 but not PC05. PC04 and PC05 activity on Oligo 1 was measured in the presence of increasing concentrations of ATP, GTP, CTP or UTP. PC04 activity towards Oligo 1 was steady up to 1.3 mM of NTPs but decreased dramatically at higher concentrations. PC05 activity, on the other hand, was not affected by the presence of any of the NTPs up to 10 mM. **(C)** The presence of trinucleotide cap analogs decreases the methylated RNA oligo product formed by PC04 and PC05. Increasing concentrations of CleanCap^®^ AG or CleanCap^®^ AU were added to PC04 or PC05 reactions containing 10 nM enzyme and 1 µM Oligo 1. The level of methylated RNA oligo decreased as the concentration of trinucleotide cap analogs increased for both PC04 and PC05. **(D)** PC04 and PC05 methylate trinucleotide cap analogs. Increasing concentrations of PC04 or PC05 were incubated with 1 µM of CleanCap^®^ AG. Both enzymes are highly active on the trinucleotide cap analog, although PC05 exhibited more robust activity than PC04 – 10 nM of PC05 was sufficient to methylate 1 µM of the cap analog at 37°C in 30 minutes whereas 40 nM of PC04 was required to achieve the same level of modification.

### One-pot two-step m6A Cap-1 mRNA IVT by pre-methylation of Cap-1 analog reagent

Intrigued by the above results, we attempted to interface enzymatic modification of trinucleotide Cap-1 analogs with IVT in a one-pot, two-step format. Cap-1 analog AG (9 mM) was incubated with buffer, PC04, or PC05 overnight before directly combining with IVT reagents (NTPs, pRNA21_AGGA DNA template, T7 RNA Polymerase, etc) (Fig 6A). A control IVT reaction using CleanCap^®^ Reagent M6 was performed to verify that the IVT conditions used supported incorporation of trinucleotide Cap-1 analogs possessing the m6A modification. The IVT product RNA was then purified for transcription yield determination and 5′ composition analysis by LC-MS/MS. We found that ∼70% (∼630 nmol in 100 µL) and ∼95% (∼850 nmol in 100 µL) of the initial 9 mM Cap-1 analog was converted to m6A Cap-1 AG in overnight PC04 and PC05 reactions, respectively (FIG 6B). IVT yield using unreacted Cap-1 analog, PC04-or PC05-modified, or CleanCap^®^ Reagent M6 were similar (1-1.5 µg/µL; FIG 6C) and of high quality (Supplementary FIG S5A). The ∼70:30 m6A Cap-1:Cap-1 analog mixture generated by PC04 resulted in a ∼40:60 m6A Cap-1:Cap-1 IVT product (FIG 6D). The low incorporation of m6A Cap-1 analog generated in this case agrees with previous reports that suggested T7 RNAP incorporates m6A Cap-1 analog less readily than Cap-1 analog (37). On the other hand, ∼95% m6A Cap-1 RNA was synthesized using the PC05-generated ∼95% m6A Cap-1 analog product (FIG 6D). RNA synthesized from the control reactions containing unreacted Cap-1 analog AG or CleanCap^®^ Reagent M6 exhibited expected methylation status and incorporation levels. The identity of the m6Am modification on the mRNA products was confirmed by nucleoside modification analysis (Supplementary FIG S5B). Therefore, in addition to the preparation of m6A Cap-1 analogs (FIG 6B), PC05 can be used to prepare millimolar concentrations of m6A Cap-1 analogs that can be directly used in IVT reactions (FIG 6D), facilitating one-pot, two-step synthesis of m6A Cap-1 RNA *in vitro*.

**Figure 6.**
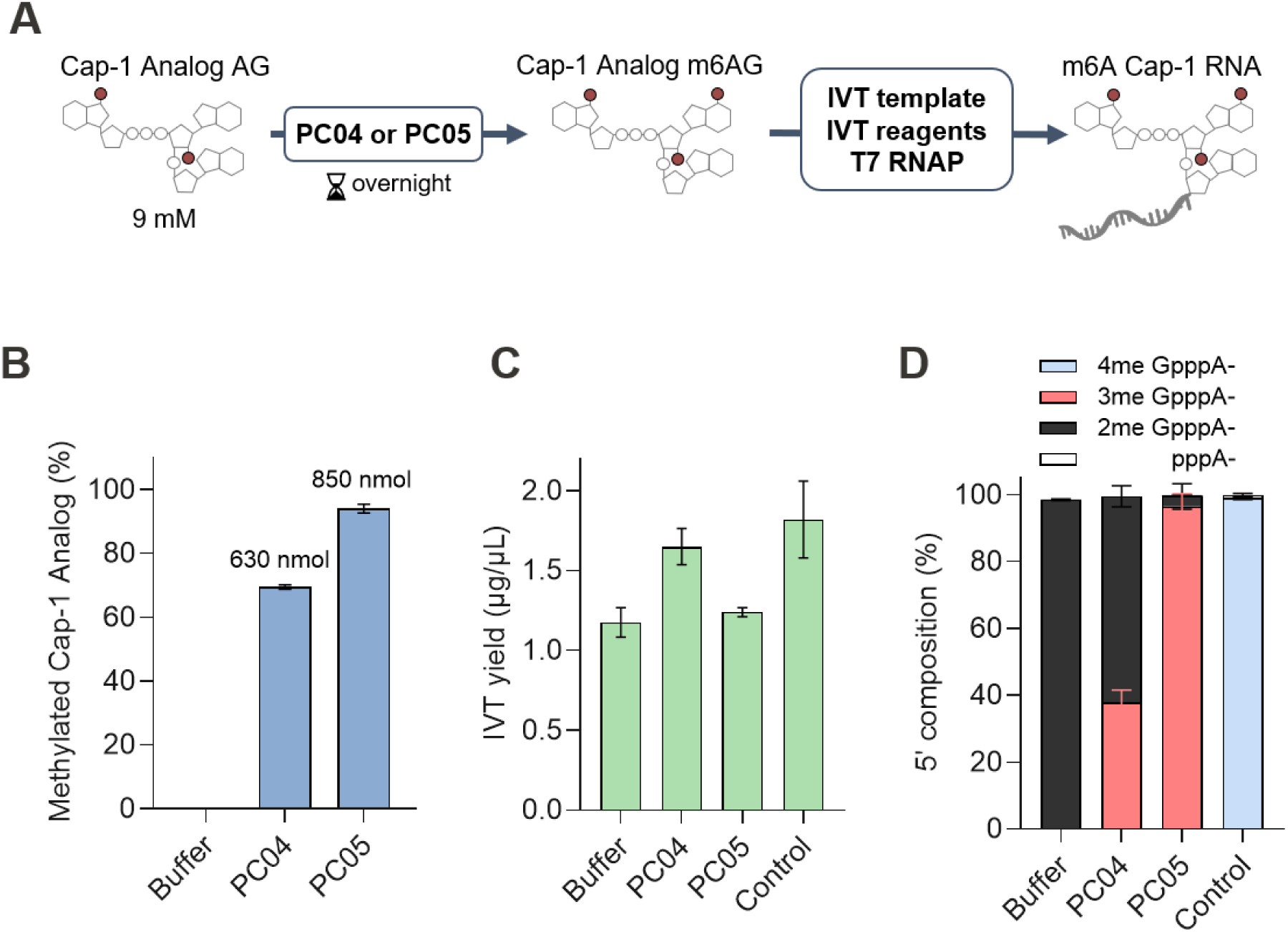
Application of PC04 and PC05 in m6A trinucleotide analog chemoenzymatic synthesis. **(A)** A two-step workflow to generate m6A Cap-1 RNA *in vitro*. Nine millimolar of CleanCap^®^ AG was incubated with 5 µM of PC04 or PC05 at 37°C overnight in the presence of 13 mM SAM. After taking an aliquot for methylation analysis (B), T7 RNA polymerase, IVT template and reagents were added directly to the PC04 or PC05 reactions which was further incubated at 37°C for 3 hours. The RNAs were purified and subjected to IVT yield and 5’ end analysis. **(B)** PC05 modified ∼95% of the cap analog (∼8.5 mM) for a total of ∼850 nmol m6A trinucleotide Cap-1 analog. PC04 modified ∼70% of the cap analog (∼6.3 mM) and generated ∼630 nmol of m6A trinucleotide Cap-1 analog. **(C)** The RNA yield of IVT using PC04-or PC05-treated trinucleotide cap analogs was comparable to IVT without PC04 or PC05 present (Buffer) or using CleanCap^®^ Reagent M6 (Control) at 1-1.5 µg/µL. **(D)** 5’ composition of IVT RNA generated using PC04-or PC05-treated trinucleotide cap analogs. IVT RNA carried ∼95% m6A Cap-1 structure (3me GpppA-) when using PC05-treated trinucleotide cap analogs. PC04-treated trinucleotide cap analog resulted in ∼40% m6A Cap-1 RNA (3me GpppA-) and ∼60% Cap-1 RNA (2me GpppA-). In the absence of PC04 and PC05 (buffer), 100% of the IVT RNA carried the Cap-1 structure (2me GpppA-). Control IVT using CleanCap^®^ M6 (3’Ome) carries a 3’O,N7-dimethyl-M6A Cap-1 structure (4me GpppA-) as expected.

### All-enzymatic one-pot m6A Cap-1 RNA synthesis

We next sought to synthesize m6A Cap-1 mRNA in a one-pot one-step format. We have previously demonstrated that Cap-0 RNA can be synthesized by using an FCE::T7RNAP fusion enzyme under standard IVT conditions (42). The Cap-0 RNA can be further modified by *Vaccinia* 2′-O MTase simultaneously in a one-pot format. Here, IVT with enzymatic co-transcriptional capping was performed using the FCE::T7RNAP fusion and *Vaccinia* 2′-O MTase in the presence or absence of 200 nM PC04 or 100 nM PC05 using the pRNA21_AGGA and ETS1 DNA templates (Fig 7B). The synthesized RNA exhibited high integrity (Supplementary FIG S6A). The presence of PC04 or PC05 did not affect the yield of IVT products (Supplementary FIG S6B). LC-MS/MS cap analysis confirmed that RNA synthesized by FCE::T7RNAP with 2’-O MTase in the absence of PCIF1 were ≥95% Cap-1 modified (Fig 7C). Reactions including PC04 generated 85% and 30% m6A Cap-1 for pRNA21_AGGA and ETS1 RNA respectively; a result consistent with biochemical experiments showing enzyme inhibition of PC04 by NTPs at the concentrations used. On the other hand, PC05 displayed high methylation activity under these conditions where m6A Cap-1 mRNA was produced to high-efficiency (≥94%) for both transcripts. It is noted that for all ETS1 IVT reactions, ∼10-20% of the RNA generated carried a 5’ diphosphate group (ppA-), indicating that the GTase activity of the FCE::T7RNAP was limiting. Quantitative nucleoside analysis confirmed the identity of the m6Am modification at stoichiometric ratios comparable to those observed by the LC-MS/MS 5′ cap analysis method (Fig 7D). These experiments demonstrate PC05 in conjunction with an RNA capping enzyme-RNA polymerase fusion and vaccinia cap 2’OMTase can efficiently synthesize m6A Cap-1 RNA *in vitro*.

**Figure 7.**
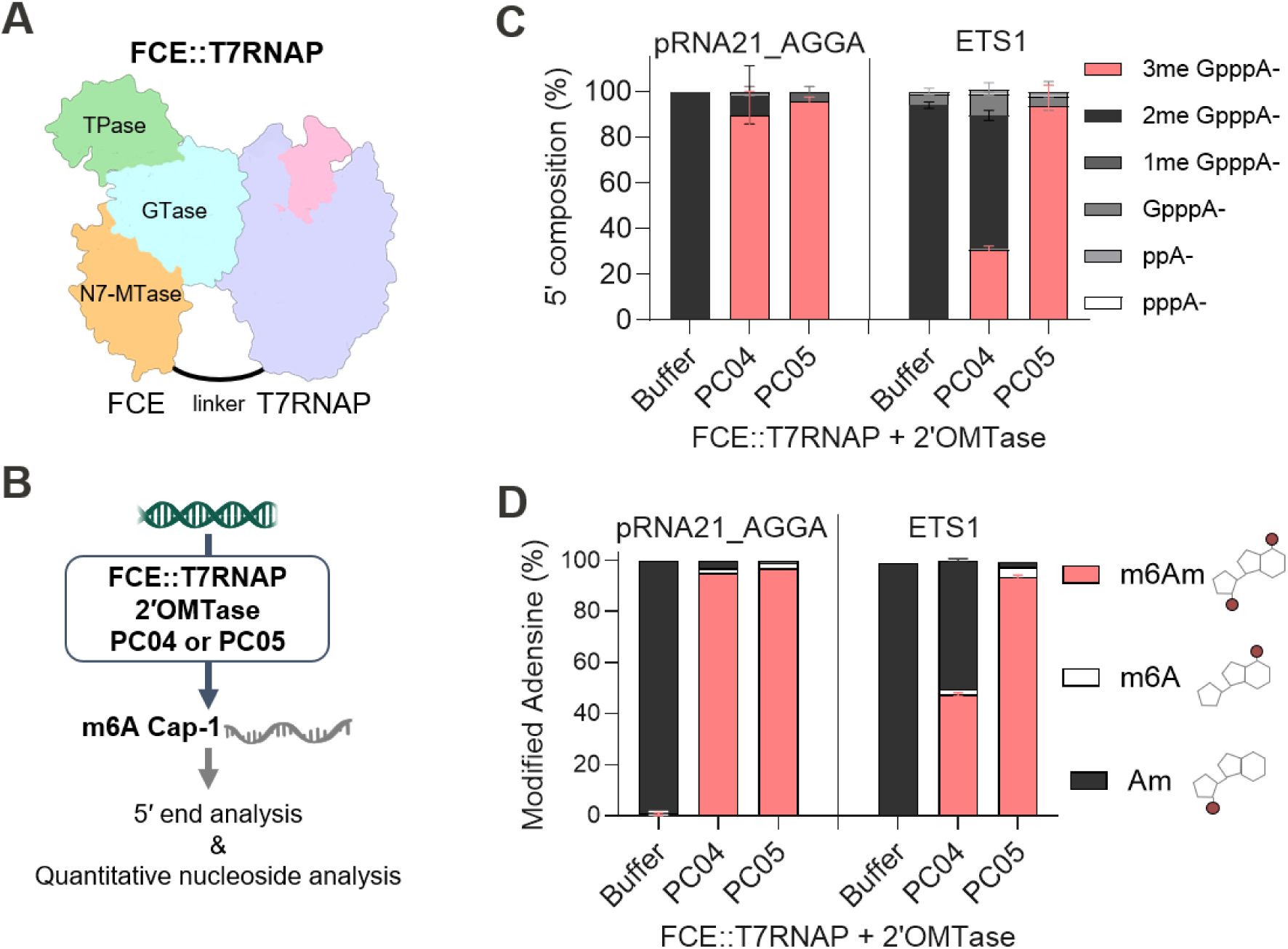
All enzymatic co-transcriptional installation of the m6A Cap-1 structure. **(A)** A schematic representation of the FCE::T7RNAP fusion (41). **(B)** One-pot all enzymatic m6A Cap-1 RNA synthesis. A linearized template was incubated with 100 mM of FCE::T7RNAP fusion, 5 U/µL of vaccinia 2’OMTase and 200 nM of PC04 or 100 nM of PC05 in the presence of 5 mM of each of the four NTPs and 0.4 mM of SAM at 37°C for 90 minutes. The RNAs were purified and subjected to RNase 4-LC-MS/MS 5’ end analysis and quantitative nucleoside analysis. **(C)** RNase 4-LC-MS/MS analysis indicated that PC04 and PC05 generated >90% m6A Cap-1 pRNA21_AGGA RNA (3me GpppA-) under the all enzymatic co-transcriptional conditions. On the other hand, while PC05 generated ∼90% m6A Cap-1 ETS1 RNA, PC04 only generated ∼30% m6A Cap-1 and ∼60% Cap-1 RNA (2me GpppA-). For all ETS1 IVT reactions, ∼10-20% of the RNA generated carried a 5’ diphosphate group (ppA-), indicating that the GTase activity of the FCE::T7RNAP was limiting. **(D)** Quantitative nucleoside analysis. The same IVT reactions were analyzed by the decapping-nucleoside analysis. The area under the peak of m6Am, m6A and Am were used to calculate the percentage of N6-adenosine methylation. The levels of m6Am detected are consistent with the intact LC-MS/MS analysis (C).

## Discussion

The m6A Cap-1 structure of mammalian messenger RNA was discovered in the 1970’s (6–8, 43) but the biological role of the modification has only begun to unfold due to the recent advances in analytical methods such as LC-MS (44) and RNA sequencing technologies (11, 45), and the discovery of PCIF1 as the sole enzyme responsible for the modification in mammalian cells (11–13, 15). While still under investigation, the biological role of m6A Cap-1 modification appears to be diverse in different cell types and may provide the cell with tunable mechanisms to regulate transcription and translation. Nonetheless, reports that synthetic mRNA containing m6A Cap-1 modification increase the duration and level of protein expression *in vivo* (36, 46, 47) suggest that the modification can be a means to improve the performance of mRNA therapeutics.

In addition to enzymatic capping, trinucleotide cap analogs are commonly used to synthesize Cap-1 RNA in IVT (48). Trinucleotide cap analogs refer to a class of synthetic cap analogs consisting of the first two nucleosides connected by a canonical 3’ to 5’ phosphate diester bond and a m7G structure connected to the first nucleotide via a 5’ to 5’ triphosphate linkage. Early kinetics studies of T7 RNA polymerase suggest that nucleotide triphosphate binding at the +2-nucleotide site of transcription precedes that at +1 site (49, 50). It has also been reported that dinucleotides pppGpA and pppGpG are potent initiators for T7 RNA polymerase *in vitro* (51, 52). It was therefore conceivable that trinucleotide cap analogs such as CleanCap^®^ AG support T7 RNA polymerase transcription initiation efficiently (48, 53).

m6A-modified trinucleotide cap analogs such as CleanCap^®^ Reagent M6 (Trilink) have been used to synthesize m6A Cap-1 RNA *in vitro*. However, addition of a methyl group to the N6 position of the initiating adenosine of these trinucleotide cap analogs requires modifications of reaction conditions such as high cap analog and T7 RNA polymerase concentrations (36, 54) or mutations in T7 RNA polymerase to facilitate efficient transcription (55). Evidence suggests that the Hoogsteen edge of guanosine and adenosine at the +1-nucleotide position is an important factor in T7 RNA polymerase transcription initiation (56, 57). For adenosine, the Hoogsteen edge consists of the N6 and N7 groups. The methyl group at the N6 position of trinucleotide Cap-1 analogs may therefore disrupt the formation of a productive initiation complex and lead to low IVT yield under standard conditions.

To overcome the mechanistic shortfall in IVT initiation, millimolar concentrations of N6A-methylated trinucleotide cap analog are needed to drive the incorporation of the cap analog into the IVT products, while only one molecule of the cap analog is required per RNA transcript. Since IVT yields oftentimes culminate in high nanomolar to low micromolar concentrations, a large majority of the cap analogs added to IVT reactions are not incorporated into the RNA transcript and discarded (57). RNA capping and cap modifying enzymes, on the other hand, turnover multiple times within the reactions using commodity co-substrates (GTP, which is already present in IVT reactions, and SAM) and are therefore a more efficient means for large scale synthesis.

Using bioinformatics and functional screening, we selected and characterized two robust homologs of human PCIF1 as a means to domesticate the cap-specific N6A methyltransferase activity and created economical and adaptable solutions to synthesize m6A Cap-1 RNA *in vitro*. The two enzymes methylate the initiating adenosine of 5′-m7G-capped RNA at nanomolar concentrations and are compatible with conditions commonly used for *in vitro* transcription and RNA capping reactions. Robust methylation activity was observed for RNA oligos, IVT transcripts containing m1Ψ, and trinucleotide cap analogs.

PC04 and PC05 can be incorporated into existing IVT workflows to synthesize m6A Cap-1 RNA efficiently (FIG 8). IVT transcripts produced using trinucleotide Cap-1 analogs or enzymatic capping can be methylated to form m6A Cap-1 after a purification step (Route 1). PC04 and PC05 can be added to FCE/VCE and Vaccinia cap 2′-O methyltransferase reactions to facilitate one-pot post-transcriptional capping and cap modification (Route 2). Taking advantage of the methylation activity on trinucleotide cap analogs, we showed that PC05 can be used to convert trinucleotide Cap-1 analogs to their m6A counterparts that can be directly used in IVT (Route 3). Ultimately, one-pot one-step m6A Cap-1 synthesis can be achieved by including PC05 in enzymatic co-transcriptional capping IVT reactions using an FCE::T7RNAP fusion protein and Vaccinia cap 2′OMTase (Route 4).

**Figure 8.**
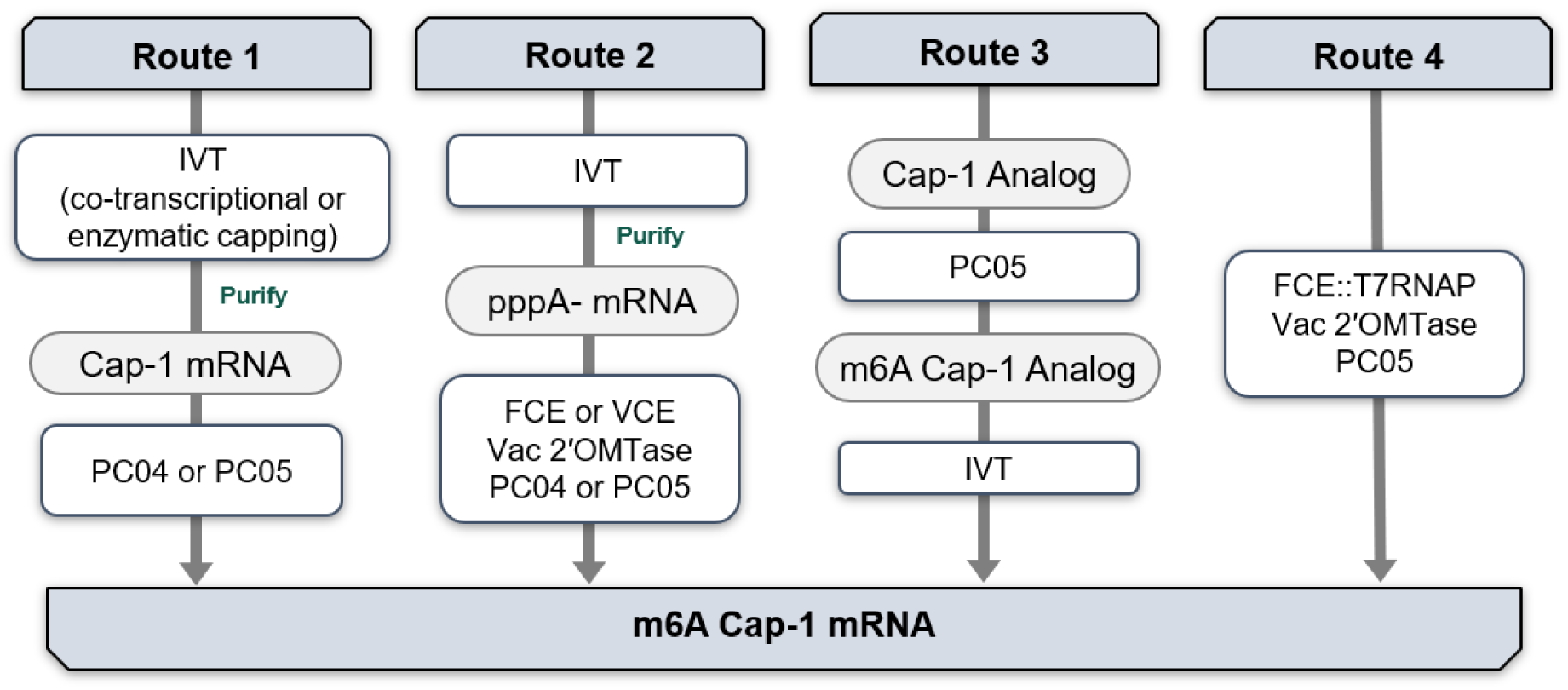
Application of PC04 and PC05 in m6A Cap-1 RNA synthesis. Route 1: Cap-1 IVT transcripts produced using trinucleotide Cap-1 analogs or post-transcriptional enzymatic capping can be methylated to form m6A Cap-1 post-transcriptionally after a purification step. Route 2: One-pot post-transcriptional 5’ capping and cap modification using PC04 or PC05 in addition to FCE/VCE and Vaccinia cap 2′-O methyltransferase. Route 3: An one-pot, two steps workflow where PC05 is used to convert millimolar concentrations of a trinucleotide Cap-1 analog to its m6A form that can be directly used in IVT reactions. Route 4: All enzymatic one-pot one-step m6A Cap-1 RNA synthesis using an FCE::T7RNAP fusion protein, Vaccinia cap 2′OMTase and PC05.

The availability of an efficient cap N6-adenosine methyltransferase such as PC04 and PC05 can enable more research on the biological role of the m6A Cap-1 modification, the establishment of more potent mRNA therapeutics and novel applications. For example, m6A Cap-1 RNA can be readily synthesized and transfected different cell types under different growth conditions to decipher their biological effects via transcriptomic analyses (63). Recombinant human PCIF1 has been used to introduce cap-adjacent adenosine with N6’-propargyl groups that allow functionalization of synthetic RNA through click-chemistry (45, 64).

In summary, we identified two RNA 5’ cap N6-adenosine methyltransferases that are highly active and can readily be incorporated into IVT workflows to produce m6A Cap-1 RNA *in vitro*. These enzymes present additional tools for research on the biological roles of the m6A Cap-1 modification and development of therapeutic mRNA.

## Methods

### Functional screening of PCIF1 candidates

PCIF1-like sequences were collected from the Joint Genome Institute Integrated Microbial Genomes & Microbiomes (65, 66). AlphaFold2 Structural predictions (39) of sequences using the Colab web server (38) were visualized using PyMol (67). Fifty-five PCIF1-like amino acids sequences were selected for functional screening. Expression plasmids (pET-29b(+)) with inserts containing *E. coli* codon-optimized genes were purchased from Twist Bioscience. A total of 250 ng of plasmid DNA was used for 30 µL *in vitro* transcription-translation reactions (IVTT) (PURExpress ΔRF123 *In vitro* Protein Synthesis Kit; New England Biolabs; E6850S) according to the manufacturer’s instructions. After a 3-hour incubation at 37°C, the solutions were added to a screening reaction mixture containing substrate RNA oligos with a 3′-desthiobiotin modification and a 5′ Cap-0 (Oligo 5, Cap0 1A 25mer[dsbtn]) or Cap-1 (Oligo 14, Cap0 1A 25mer[dsbtn]) structure. The 50 µL reactions containing 5% (v/v) IVTT solution, 0.6 µM RNA, 1 U/µL Murine RNase Inhibiter and 0.2 mM SAM in 1x capping buffer (New England Biolabs, B2080AAVIAL) were incubated at 37°C for 1 hour. The RNA oligo was then purified using hydrophilic streptavidin-coated magnetic beads (New England Biolabs; S1421S). Reactions were mixed with the beads at a 1:1 ratio and incubatedfor 30 min at room temperature before being washed thrice with a wash buffer (5 mM Tris, pH 7.5, 60 mM NaCl, 0.5 mM EDTA). RNA was eluted in 30 µL nuclease-free water by heating to 65°C for 4 minutes. Samples were filtered in 0.22 µm centrifugal filters (Millipore; UFC30GV0S) and methylation status was analyzed by intact oligo LC-MS/MS. Plasmids encoding Vaccinia VP39 (a Cap-0 2′OMTase) served as a positive control for IVTT and cap modification assay.

### IVT Template plasmid construction

IVT templates were designed to encode two 2.1 kb RNA molecules each containing the first 50 bases of a 5′UTR, the consensus mammalian Kozak sequence, a firefly luciferase coding sequence, a 3′UTR derived from the publicly available sequence of BNT162b2, a 60-adenosine tail, and a BspQI restriction site. Transcription of the RNA is driven by the T7 Phi2.5 promoter. The 5’UTRs were a derivation of pRNA21 with a 5’-AGGA extension (pRNA21_AGGA) or the human ETS1 mRNA (ETS1; NCBI ID NM_001143820.2). The DNA sequences of the T7 transcription template are available in Supplementary Table S2. The cloning of the inserts into pUC57, preparation and linearization of the plasmid DNA were done commercially (Genscript). Supplementary Table S3 shows the expected RNA transcript sequences based on the DNA templates.

### PC04 and PC05 expression and purification

Protein expression and purification for data presented here was performed commercially (Genscript). Expression constructs containing the PC04 and PC05 ORFs with a C-term 6x His-Tag were inserted into pET29b(+) and used to transform *E. coli* T7Express *lysY/I^q^* (New England Biolabs, C3013I). Recombinant protein expression was induced by 0.5 mM IPTG once cell density reached OD_600nm_=0.6-0.8 and incubation at 15°C for 16 hrs. The proteins were purified from cell lysate supernatant in two steps using a Ni-NTA column and size exclusion chromatography (Superdex 200). Protein purity was >90% by Comassie blue-stained SDS-polyacrylamide gel. Stocks of each enzyme were prepared by dialyzing into a storage buffer containing 50% glycerol. Working stocks of 10 µM enzyme, quantified using the Bradford colorimetric assay (Bio-Rad), were stored at -20°C for over 6 months with no observable loss of activity.

### Identification of m6A Cap-1 modification

Nucleoside analysis was performed to verify the identity of the methylation products of the PCIF1-like enzymes (FIG 2D). Oligo 14, a synthetic 25mer with the Cap-1 structure (m7GpppAm-) containing a single cap-adjacent 2’-O-methyl-adenosine was used such that the presence of the N6,2′-O-dimethyladenosine (m^6^Am) is indicative of cap-specific N6-adenosine methyltransferase activity after decapping. The methylation reaction containing 2 µM of enzyme and 1.5 µM of substrate was incubated at 37°C for 2.5 hours. Since no validated m6A Cap-1 dinucleotide (m7Gpppm6Am) standard substance is commercially available, the methylation reaction products were decapped before subjecting the RNA to nucleoside digestion such that the presence of 2’-O,N6-dimethyladenosine (m6Am) is indicative of the formation of m6A Cap-1. Ten micrograms of reaction products were decapped using mRNA decapping enzyme (New England Biolabs; M0608S) and purified using hydrophilic streptavidin-coated magnetic beads (New England Biolabs; S1421S) yielding RNA with monophosphate (p-) 5′ ends. The oligos were converted to nucleosides using Nucleoside Digestion Mix (New England Biolabs; M0649S) and filtered using 0.22 µM spin column (Millipore; UFC30GV25). The modified nucleotide species present in each sample was resolved using UHPLC-MS/MS analysis with dynamic multiple reaction monitoring with technical duplicates (dMRM).

### Enzyme activity assays using RNA oligomer substrates

Enzyme activity assays were conducted in 1x capping buffer (New England Biolabs, B2080AAVIAL) containing 400 µM SAM, 1 U/uL Murine RNase Inhibiter and indicated concentrations of RNA oligos in duplicate unless specified otherwise. Reactions at 30 µL final volume were incubated at 37°C for 30 minutes and stopped by heating to 75°C for 5 minutes in a thermocycler. Enzymes stocks were diluted in 1x capping buffer for assays unless specified. A 3′-desthiobiotin TEG modification ([dsbtn]) is present in indicated RNA substrates to enable purification of the oligos from the reaction mixtures for compatibility with LC-MS/MS. Percent composition of methylated oligos was determined for each sample by mass signal intensity of the corresponding species. The mean ± 1 standard deviation of duplicate experiments are reported. A table of oligo substrates is found in Supplementary Table 1.

Substrate specificity experiments presented in FIG 2C were performed on RNA oligos with different 5′ chemical groups (oligos **2-10** in Supplementary Table S1) using 10 nM enzyme and 1.5 µM substrate. Oligo 3, which contains a 5’ diphosphate group was generated by incubating oligo 2 (containing a 5’ triphosphate group) with FCE (NEB) in the absence of co-substrates GTP or SAM. Oligo 4 contains an unmethylated G-cap generated by incubating oligo 2 with FCE in the presence of GTP. Oligo 5 contains a Cap-0 structure generated by incubating Oligo 2 with FCE, GTP and SAM. The reacted oligos were purified using hydrophilic streptavidin-coated magnetic beads (New England Biolabs; S1421S).

Enzyme titration assays (FIG 3A, D) were performed by incubating serial dilutions of PC04 or PC05 with 1µM of Cap-1 RNA oligo substrates in 1x reaction buffer at 37°C for 30 minutes, followed by heating to 75°C for 5 minutes. Oligo 1 (Cap-1 1A 12mer[dsbtn]) is a 12mer with a single Cap-adjacent 2′-O-methyladenosine. Oligo 6 (Cap-1 8A 25mer[dsbtn]) is a 25mer containing 8 internal adenosines. Oligos 12 (Cap-1 AU 12mer) and 13 (Cap-1 Am1Ψ 12mer) are 12mers designed specifically to place a uridine or N1-methylpseudouridine at position 2, adjacent to the methyl acceptor 2′-O-methyladenosine.

To investigate the temperature range of the enzymes (FIG 3B), reaction mixtures were set up in pre-chilled tubes in 4°C cold blocks to ensure synchronized initiation. Reaction mixture (270 µL) containing 2 nM of enzyme and 1 µM oligo 1 (Cap-1 1A 12mer[dsbtn]) were aliquoted and placed at temperatures ranging from 15-65°C in a thermocycler with variable heating zones (Applied Biosystems VeritiPro) to initiate the reactions.

To investigate the pH range of the enzymes (FIG 3C, Supplementary FIG S4), a final concentration of 50 mM Tris, HEPES, Bis-Tris, or Bis-Tris Propane solutions (Hampton Research) at the indicated pHs were added to a minimal reaction buffer (400 µM SAM, 2.5 mM Tris pH 7.0, 5 mM KCl, 1 mM MgCl_2_, 1 mM DTT, 1 U/uL Murine RNase Inhibiter, pH 7.0 at 25°C) and 1 µM of oligo 1 (Cap-1 1A 12mer[dsbtn]). Reactions were initiated with the addition of enzyme diluted in minimal reaction buffer at a final concentration of 2.5 nM.

For RNA methylation activity assays in the presence of NTPs or Cap-1 analogs (FIG 5B, C), 0.2-10 mM of each of the individual NTPs or CleanCap^®^ Reagent AG (Trilink; N-7113) or CleanCap^®^ Reagent AU (Trilink; N-7114) were incubated with 5 nM enzyme and 1 µM oligo 1 (Cap-1 1A 12mer[dsbtn]) substrate at 37°C for 30 min, followed by heat inactivation and LC-MS/MS analysis as indicated above.

### N6A-methyation reactions on purified synthetic mRNA

Substrate RNA was made by IVT using HiScribe T7 High Yield RNA Synthesis Kits (New England Biolabs; E2040S) with or without trinucleotide Cap-1 analogs following manufacturer’s recommended protocols. N1-methylpseudouridine modified RNA was made by substituting UTP with m1ΨTP. After IVT, the DNA template was removed by adding 1 µL of DNase I-XT (New England Biolabs; M0570S) per 20 µL of IVT and incubating at 37°C for 20 min. The RNA was purified using Monarch RNA Cleanup kits (New England Biolabs; T2040S) according to the recommended protocol. For enzymatically capped RNA, purified uncapped IVT RNA was capped to Cap-1 using FCE with Vaccinia 2’OMTase (New England Biolabs; M0366S) according to recommended protocol and purified. RNA quantification was done using a NanoDrop instrument (Thermo Fisher). Size and integrity were assessed by RNA Tapestation and RNA ScreenTapes (Agilent). For each duplicated experiment, a 5′ cap composition analysis was performed using the RNase 4-LC-MS/MS method detailed in (41) and described below. Reaction volumes varied from 50 to 80 µL to ensure a minimum of 40 µg of mRNA for cap analysis. The percent composition of methylated oligos was determined for each sample by MS signal intensity of annotated fragments. The mean ±1 standard deviation of replicated experiments are reported.

Purified Cap-1 RNA at a concentration of 0.75 µg/µL (∼1 µM) that had been enzymatically capped or co-transcriptionally capped with a Cap-1 analog was allowed to react with 120 nM of PC04 or PC05 in 1x capping buffer (New England Biolabs, B2080AAVIAL) with 1 U/µL Murine RNase Inhibiter) containing 400 µM SAM at 37°C for 60 min.

One-pot, three-enzyme m6A Cap-1 capping on purified uncapped IVT RNA was performed in reactions containing 400 µM SAM, 500 µM GTP, 120 nM PC04 or PC05, 5 U/µL Vac 2’OMTase and 1 U/µL VCE or FCE in 1x capping buffer (New England Biolabs, B2080AAVIAL) containing 1 U/uL Murine RNase Inhibiter. The VCE reactions contained 0.5 µg/µL substrate RNA. The FCE reactions contained 0.75 µg/µL substrate.

### RNA 5′ group analysis

The composition of mRNA 5′ end was determined by RNase 4 endonuclease protection assay published previously (41). Briefly, an antisense DNA oligo probe containing a desthiobiotin modification is annealed to the 5′ 18-25 bases of an mRNA and digested with RNase 4 (New England Biolabs; M1284S) using 1 µL of 50U/µL RNase 4 in 1x NEB Buffer r1.1 (New England Biolabs; B6001SVIAL). Hybrid duplexes containing the DNA probe and the protected 5′ mRNA fragments are then purified and enriched using hydrophilic streptavidin-coated magnetic beads (New England Biolabs; S1421S). The fragments are subjected to LC-MS/MS analysis. The mass of RNase 4 cleavage fragments matching the relevant 5′ structures (pppA-, ppA-, GpppA- and GpppA-with 1, 2 or 3 methyl groups) were assigned to the appropriate 5′ structures. Specifically, masses that are consistent with GpppA-plus one, two and three methyl groups are assigned Cap-0, Cap-1 and m6A Cap-1 respectively. The mass intensities of the assigned RNase 4 cleavage fragments were used to estimate the percentage of the corresponding 5’ structure in the mRNA preparations.

To verify the identity of m6A Cap-1 on RNA, the methylation reaction product RNA was decapped using mRNA decapping enzyme (New England Biolabs) according to manufacturer’s instructions. The decapped mRNA was purified using Monarch RNA Cleanup kit (New England Biolabs; T2040S). The purified decapped mRNA was incubated overnight with nucleoside digestion mix (New England Biolabs; M0649S) and subjected to nucleoside analysis using LC-MS/MS with dynamic multiple reaction monitoring (dMRM). With decapping, m6Am detected in LC-MS/MS is attributed to the N6-adenosine methyltransferase activity of the PCIF1 enzymes and 2’-O-methyltransferase activity of vaccinia cap 2’-O-methyltransferase activity. The mass intensity of the detected m6Am, m6A and Am was used to estimate the level of m6A Cap-1 incorporation.

### Trinucleotide Cap-1 Analog methylation assays

For activity assays using Cap-1 analog as substrates, 1 µM CleanCap^®^ Reagent AG (Trilink; N-7113) or CleanCap^®^ Reagent AU (Trilink; N-7114) was incubated with 0.6 to 40 nM of PC04 or PC05 in 1x capping buffer (New England Biolabs, B2080AAVIAL) at 37°C for 30 min in duplicate. The reactions were stopped by heating at 80°C for 5 min and the mass composition of methylated CleanCap^®^ Reagent AG was determined by LC-MS/MS.

High trinucleotide cap analog concentration reactions (FIG 6A) combined 4 or 8 mM of CleanCap^®^ Reagent AG (Trilink; N-7113) with 5 µM PC04 or PC05 enzyme in a reaction buffer (50 mM Bis-Tris, 100 mM NaCl, 1 mM DTT, 5% DMSO, pH 6.8 at 25°C). Methyl doner and MgCl_2_ concentrations were adjusted according to the substrate concentration. Reactions containing 4 mM substrate had 8 mM SAM and 4 mM MgCl_2_ while 8 mM substrate reactions had 12 mM SAM and 8 mM MgCl_2_. Reactions were incubated at 37°C for 18 hours and stopped by heating at 80°C for 5 min. Composition of the methylated CleanCap^®^ Reagent AG was determined by LC-MS/MS. The mean ±1 standard deviation of quadruplicate experiments is reported.

### One-pot two-step m6A Cap-1 synthesis using trinucleotide Cap-1 Analog

For reactions combining enzymatic methylation activity on Cap-1 analog with IVT (FIG 6B-F), 100 µL reaction mixtures of 9 mM CleanCap^®^ Reagent AG (Trilink; N-7113) and 5 µM PC04 or PC05 in a reaction buffer (50 mM Bis-Tris, 5 mM NaCl, 1 mM DTT, 5% DMSO, pH 6.8 at 25°C) containing 13 mM SAM and 9 mM MgCl_2_ were incubated for 18 hours at 37°C. The reaction was stopped by heating to 80°C for 5 min and an aliquot was removed for product analysis by LC-MS/MS. The overnight reactions were mixed with a solution containing IVT reagents at a 3:1 ratio to achieve final conditions of 6 mM Cap analog and 3 mM NTPs. The 120 µL IVT reaction conditions were as follows: 50 mM Bis-Tris pH 6.8, 5 mM DTT, 15 mM MgCl_2_, 2.12 mM spermidine, 1 U/µL Murine RNase Inhibitor, 3 mM NTP mix, 1 U/mL *E. coli* Inorganic Pyrophosphatase, 0.05 µg/µL DNA template (pRNA21_AGGA), and 9375 U/mL T7 RNA Polymerase (New England Biolabs; M0251S). A solution of 9 mM CleanCap^®^ Reagent M6 (CleanCap^®^ Reagent m6AG (3’Ome); Trilink; N-7453), 50 mM Bis-Tris pH 6.8, 5 mM NaCl, 1 mM DTT, 9 mM MgCl_2_, and 5% DMSO was used in place of the PCIF1 reactions as a positive control for IVT with co-transcriptional capping using an m6A modified Cap-1 analog. The solutions were incubated at 37°C for 3 hours with gentle inversion at 30 min intervals. DNA template was removed by adding 6 µL of DNase I-XT (New England Biolabs; M0570S) for 37°C for 20 min. The RNA was purified by mixing with RNACleanXP magnetic beads (Beckman Coulter; A63987) at a 1:1.8 volume ratio, washed three times with 70% ethanol, and eluted in 1 mM sodium citrate, pH 6.5. The size and integrity of the purified RNA was assessed by Tapestation and RNA ScreenTape (Agilient) (Supplementary FIG S6A). RNA 5′ analysis was performed using the RNase 4-LC-MS/MS method. The identity of the added m6A Cap-1 structure was verified using the nucleoside analysis workflow described above.

### One-pot all-enzymatic m6A Cap-1 IVT

For one-pot m6A Cap-1 synthesis using the FCE::T7RNAP fusion enzyme (FIG 7), 100 µL IVT reactions containing 200 nM PC04 or 100 nM PC05 were mixed with 0.05 µg/uL DNA template, 100 nM FCE::T7RNAP, 5 U/µL Vac 2′OMTase, 5 mM of each NTP, 400 µM SAM, 1U/µL Murine RNase Inhibitor, 5 U/µL *E. coli* pyrophosphatase, 5 mM DTT, an additional 14 mM MgCl_2_, in 1x T7 RNAP Polymerase Reaction Buffer (40 mM Tris-HCl, 6 mM MgCl_2_, 1 mM DTT, 2 mM spermidine, pH 7.9 at 25°C). Reactions were incubated at 37°C for 90 minutes. DNA template was removed by adding 5 µL DNase I-XT (NEB) and incubating at 37°C for 20 min. The IVT products were purified using Monarch RNA Cleanup kits (New England Biolabs; T2040S) according to the manufacturer’s instructions. The IVT yield was assessed using a NanoDrop instrument (Thermo Fisher). RNA 5′ analysis was performed using the RNase 4-LC-MS/MS method. Decapping-nucleoside analysis was used to confirm the identity of modified nucleosides.

## Acknowledgments

We thank Brett Robb and Ivan Corrêa for support and direction; Christa Molé, Eric Wolf, Albert Tsai, and Sean Johnson for helpful discussions, and facilities and administrative personnel for daily support of the operation.

## Funding and additional information

This work is funded by New England Biolabs, Inc.

---

**FIGURE S1.**
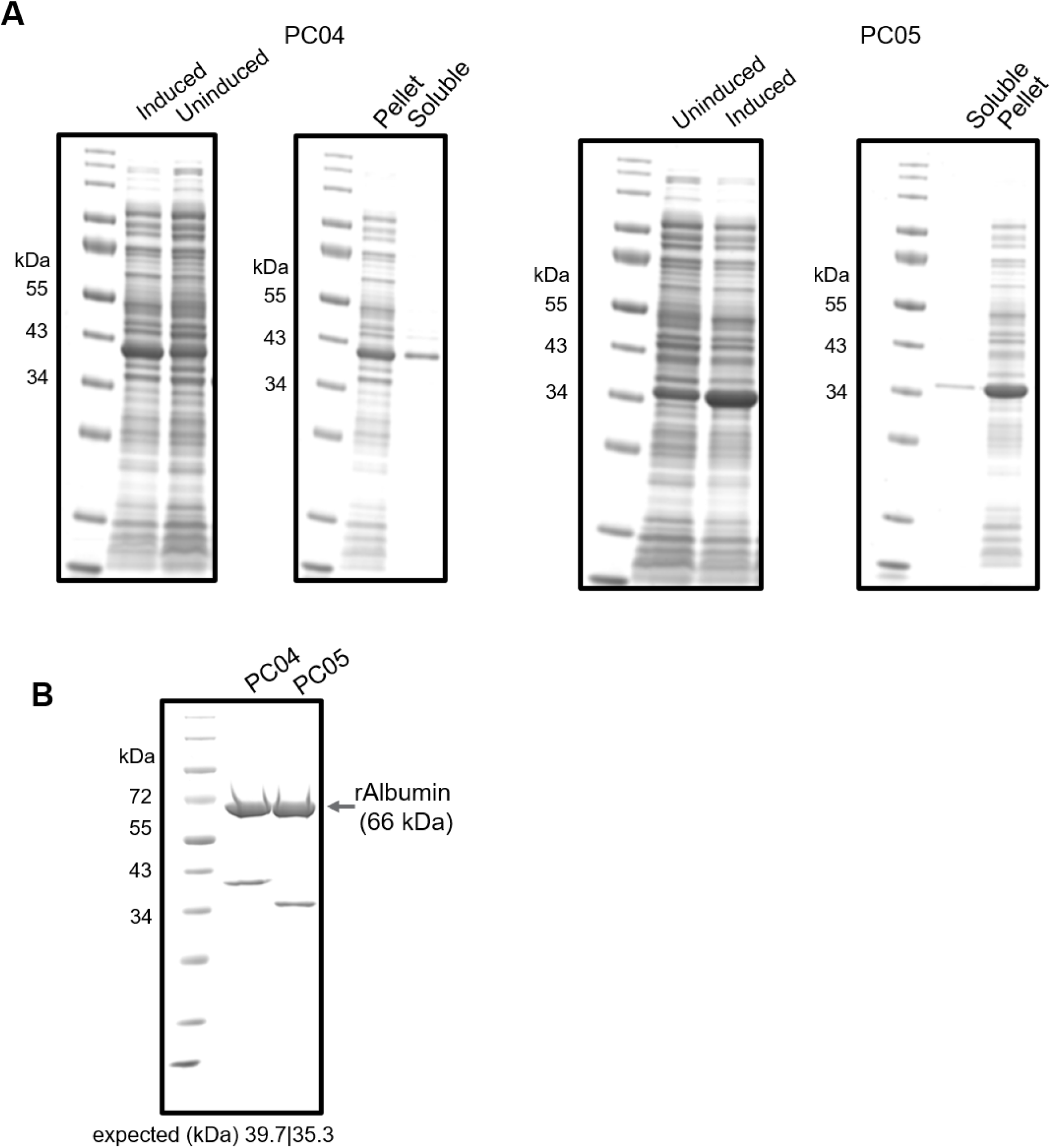
PC04 and PC05 expression and purification. A) SDS-PAGE of recombinant PC04 and PC05 *E. coli* expression and lysate. B) SDS-PAGE of purified PC04 and PC05. Enzyme stocks were diluted in buffer containing recombinant albumin as internal control for protein quantitation.

**FIGURE S2.**
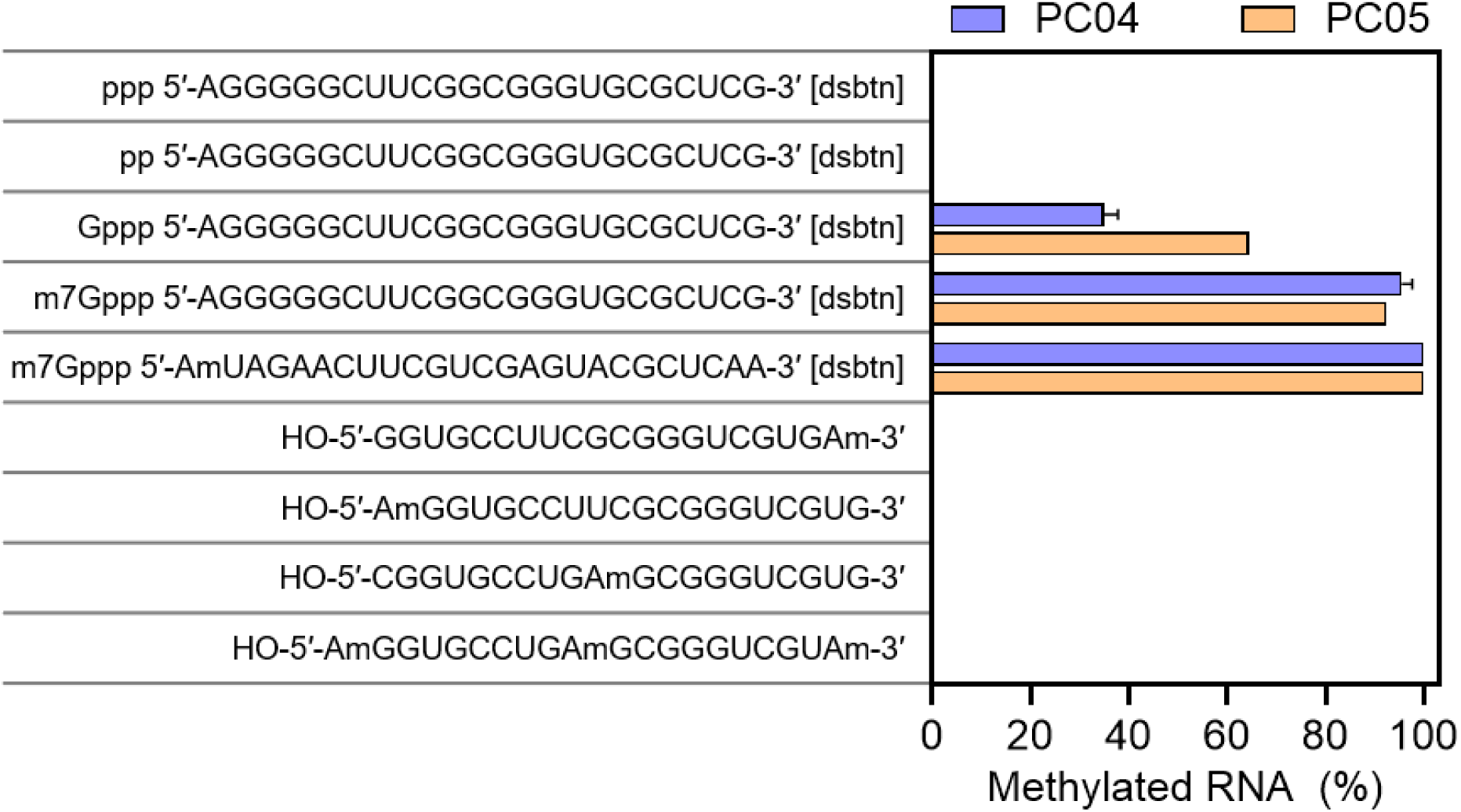
Expanded results for substrate specificity. PC04 and PC05 at 10 nM was allowed to react with 1.5 µM of RNA oligonucleotides with different sequences and modifications (Oligo index 2-10) for 30 minutes at 37°C. Methylation of each oligo was monitored by LC-MS/MS and product composition was plotted. Datapoints are average values from duplicated experiments. Error bars represent one standard deviation.

**FIGURE S3.**
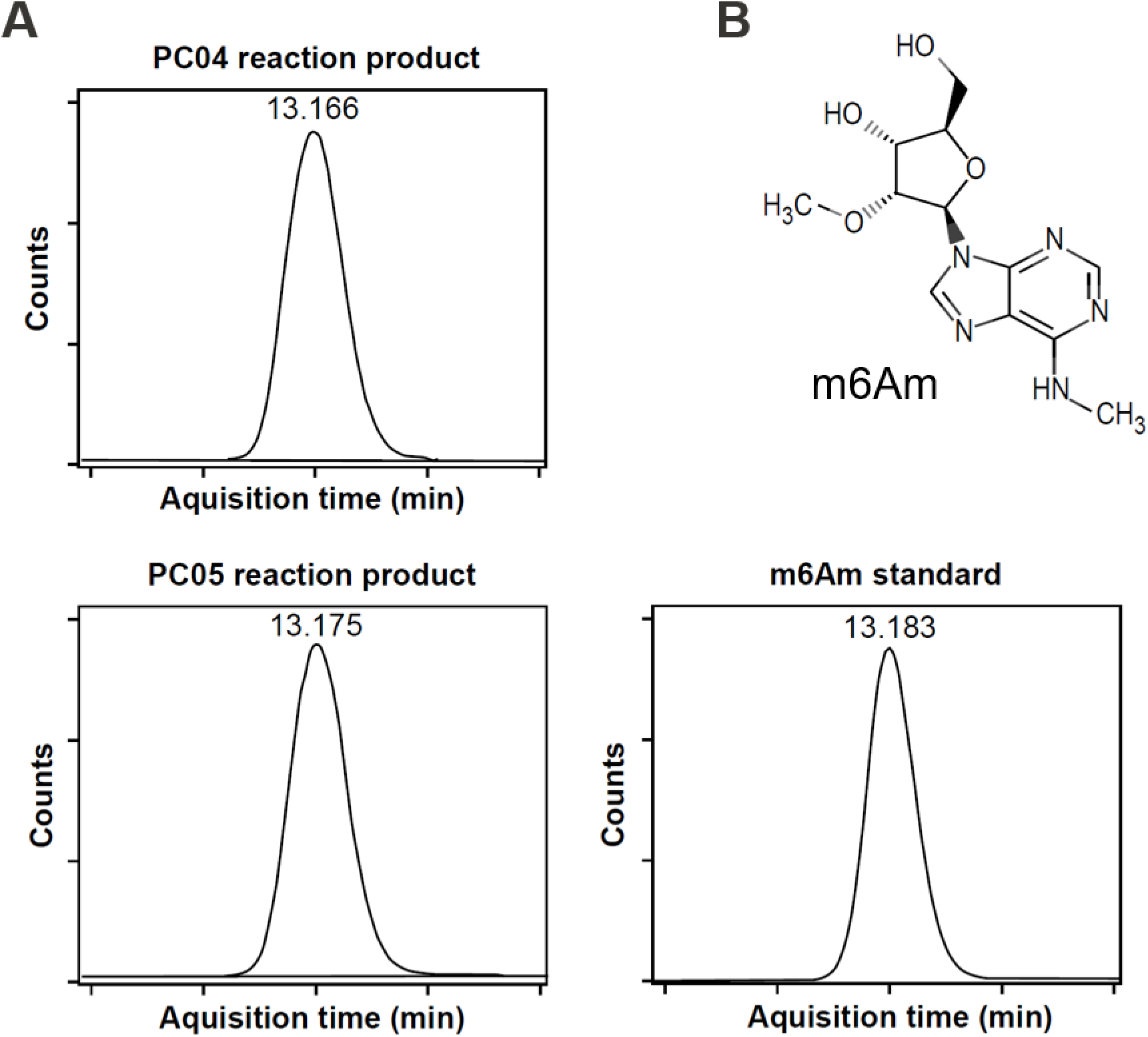
Verification of PC04 and PC05 methylation products. **(A**) Retention time and dynamic multiple reaction monitoring. After purification, decapping and digestion into nucleosides, the PC04 or PC05 reaction products were subjected to UHPLC-MS/MS analysis with dynamic multiple reaction monitoring (dMRM). The retention time of the major reaction product is 13.166 min and 13.175 min for PC04 and PC05, respectively, compared to that of 13.183 min of the N6-methyl-2’-O-methyladenosine (m6Am) standard. The mass of the PC04 and PC05 main products transitioned from 296.1 Da (consistence with the mass of m6Am) to 150.1 Da (consistent with the mass of N6-methyladenine), further supporting the notion that PC04 and PC05 produced the m6Am cap. The UHPLC trace of no enzyme control has been overlayed to the PC04 and PC05 traces (grey lines). **(B)** Chemical structure of m6Am.

**FIGURE S4.**
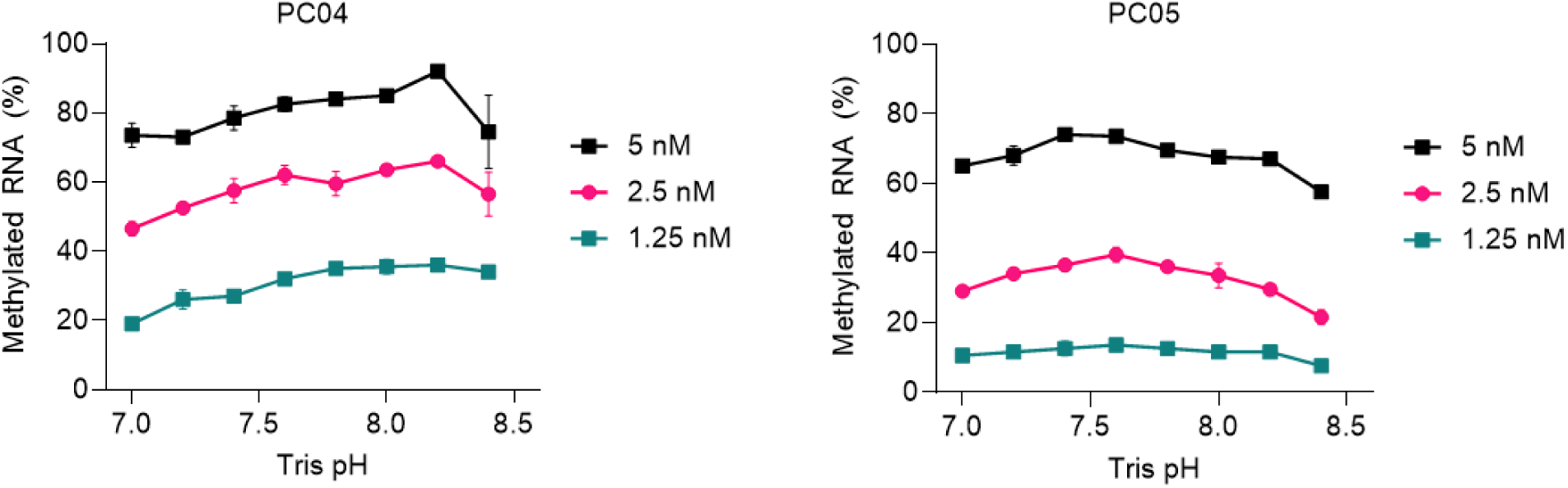
PC04 and PC05 activity as a function of the pH value of Tris buffer. PC04 or PC05 (5, 2.5 and 1.25 nM) was incubated with 1 µM of Cap-1 RNA oligo (Oligo 1) in the presence of 50 mM Tris at different pHs. Datapoints are average values from duplicated experiments. Error bars represent one standard deviation.

**FIGURE S5.**
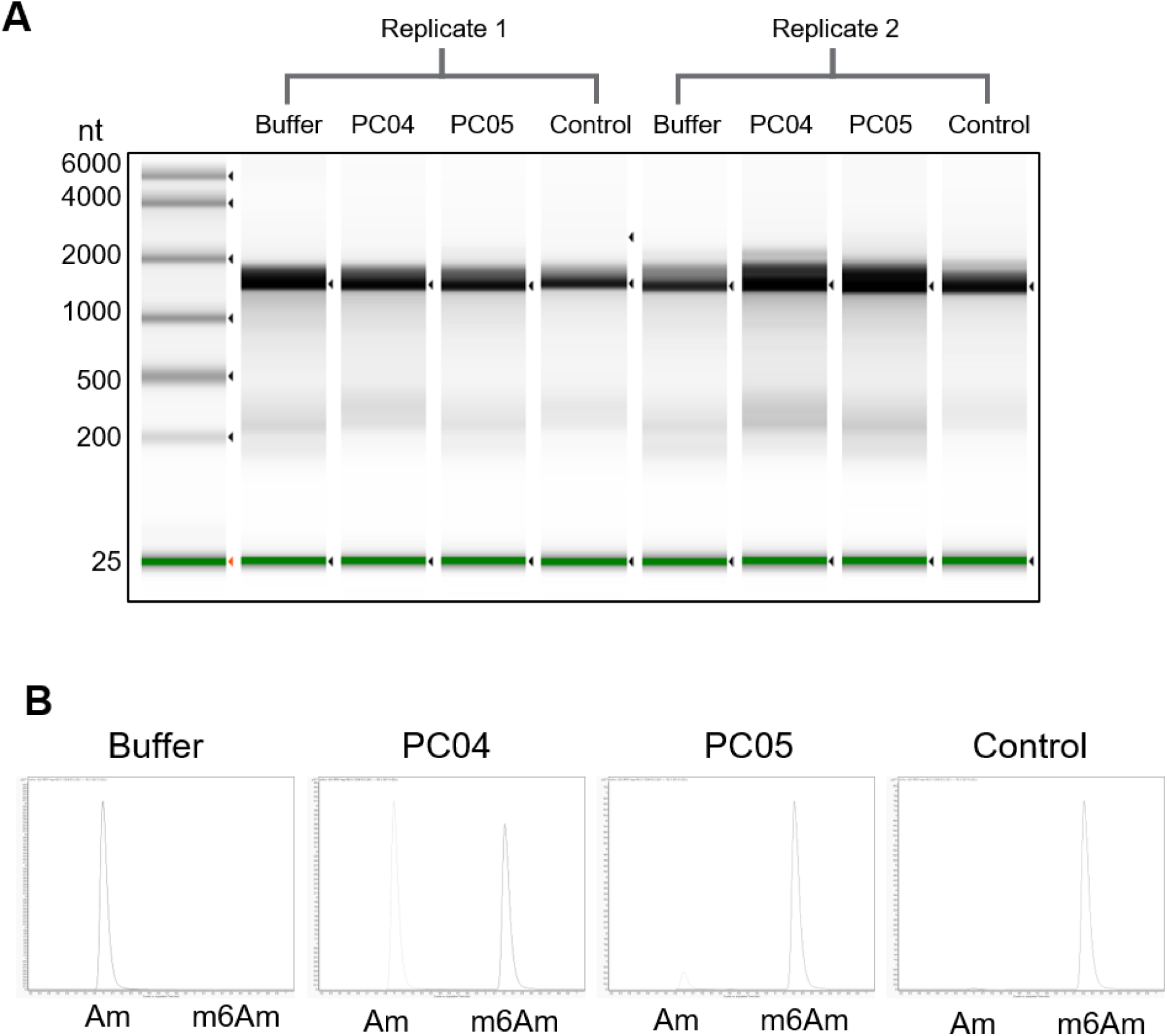
RNA analysis from interfacing Cap-1 Analog modification with IVT. **(A)** Tapestation analysis of IVT RNA after purification. **(B)** Representative dMRM nucleoside analysis of the 5’ group of the IVT RNA.

**FIGURE S6.**
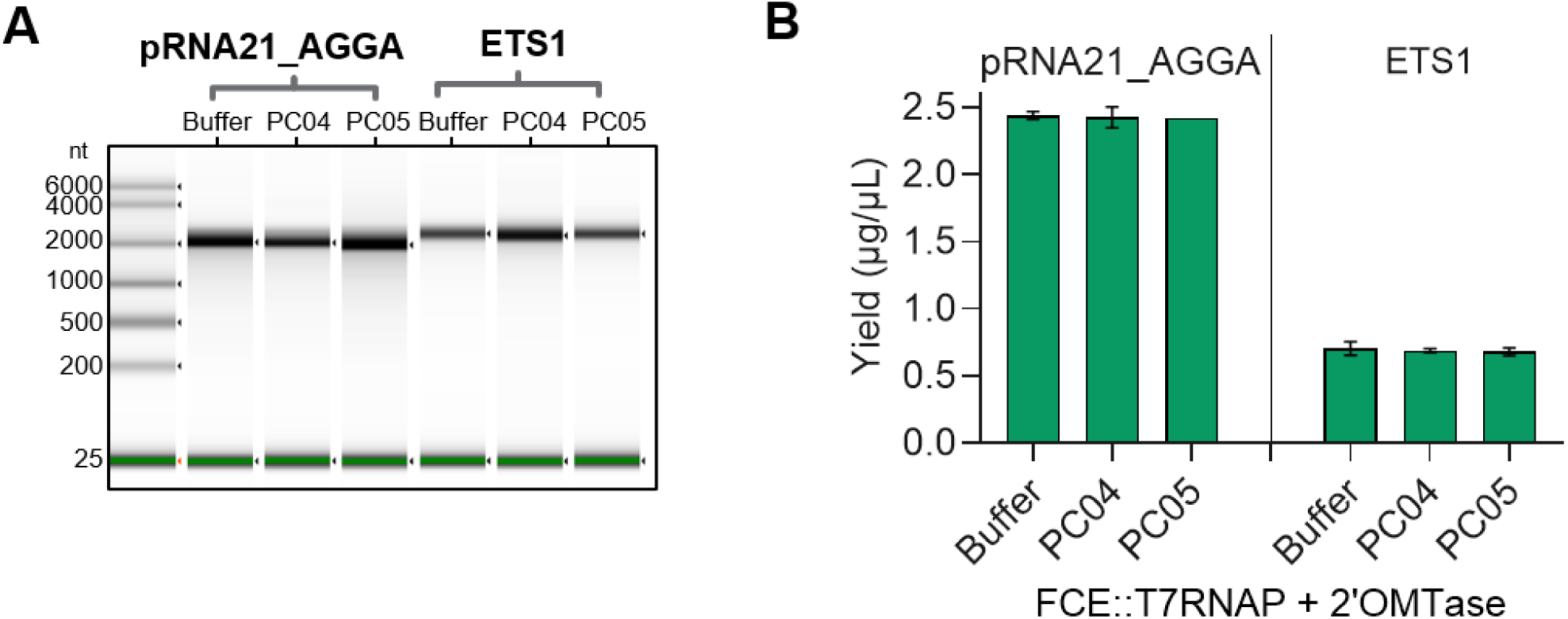
RNA analysis of RNA generated by co-transcriptional capping using FCE::T7RNAP. (A) Representative integrity analysis of RNA material using Tapestation. **(B)** IVT yield of purified RNA product determined by absorbance at 260 using Nanodrop. Datapoints are average values from duplicated IVT experiments. Error bars represent one standard deviation.

**Table S1.**
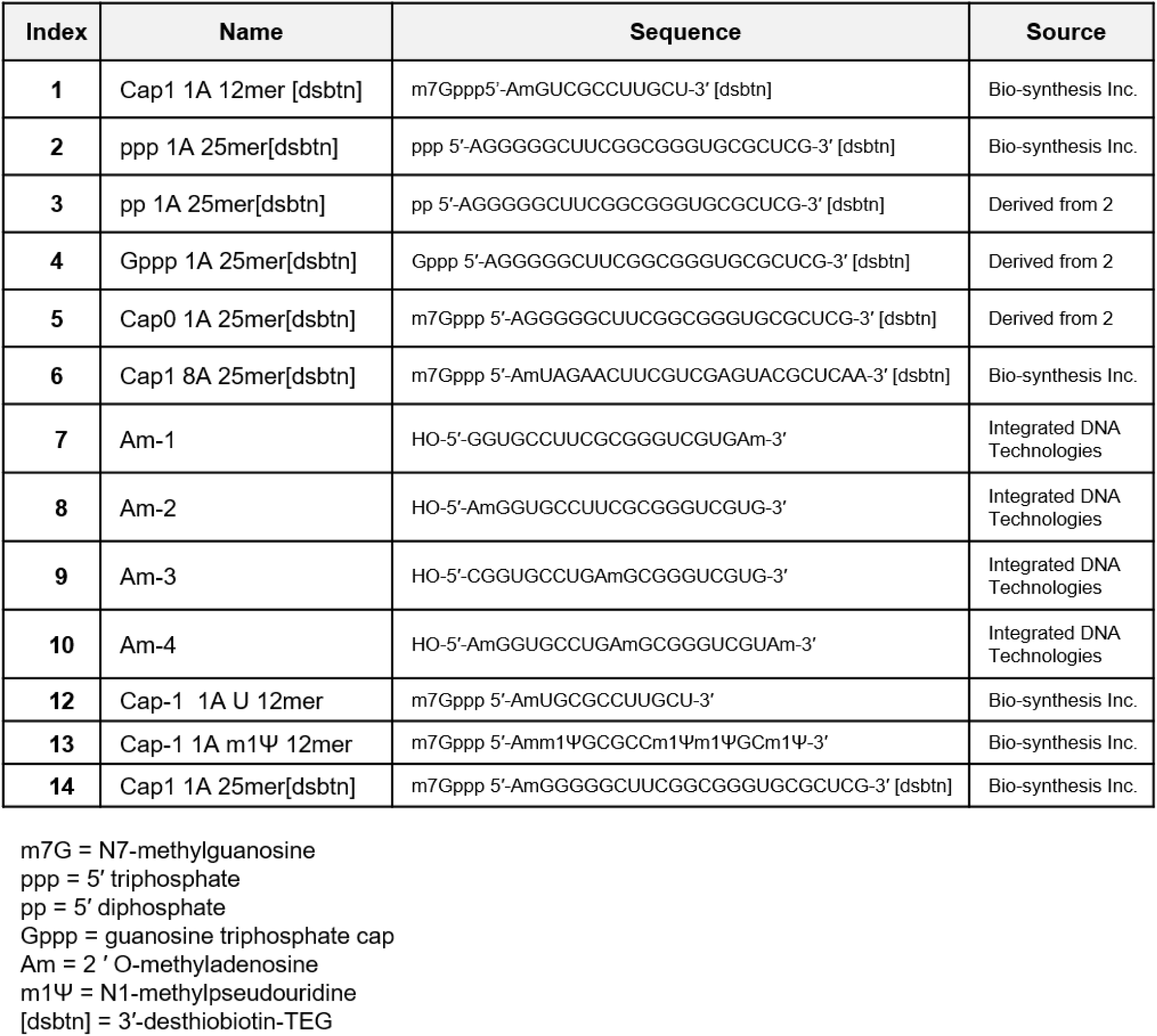
RNA oligos used in this study. RNA Oligos referred to by index number or name. Capped RNA Oligos were ordered commercially synthesized or oligo 2 was capped using Faustovirus Capping Enzyme (FCE) and/or *Vaccinia* 2’O-methytransferase with conditions.

**Table S2.**
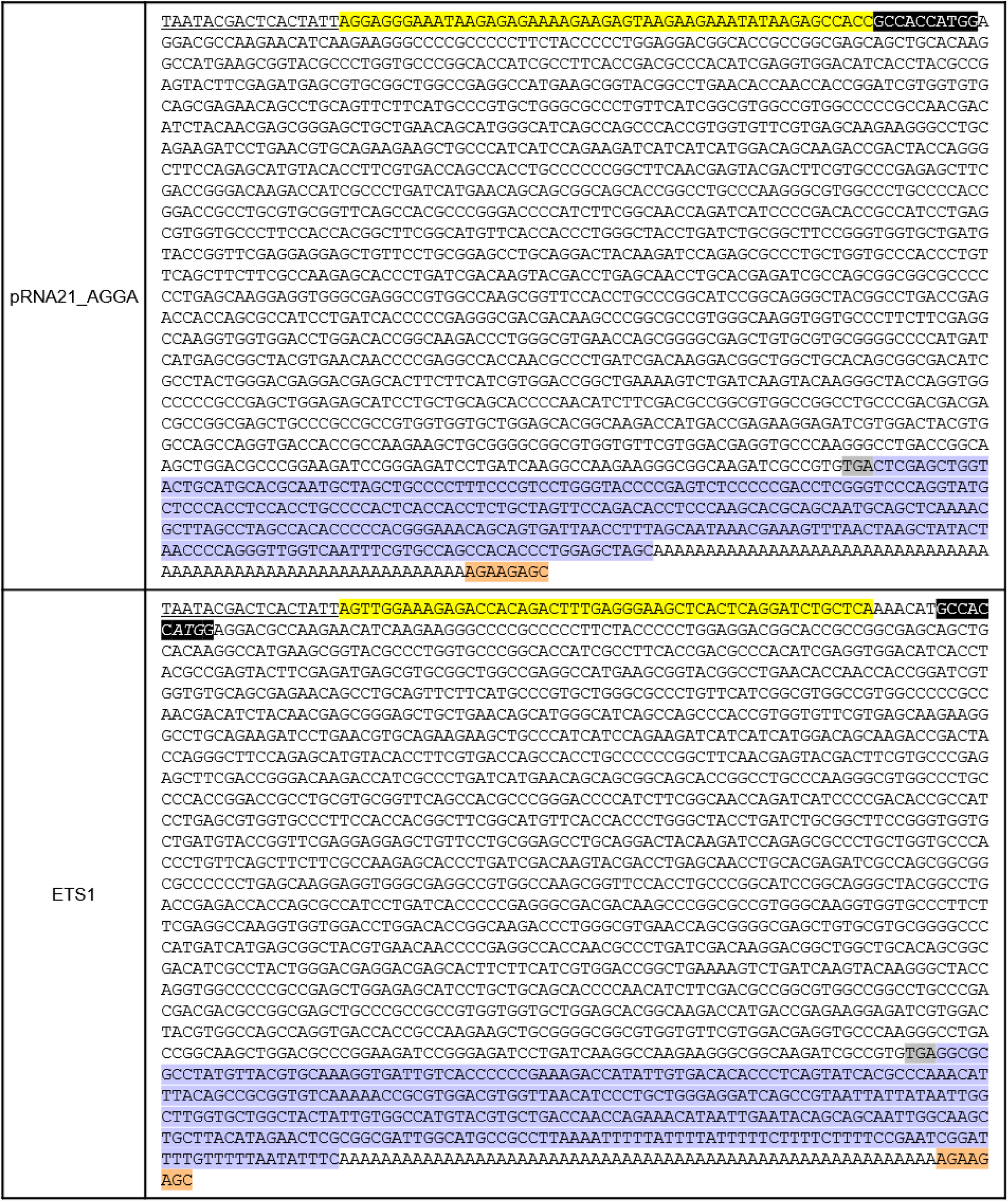
IVT template sequences of the model mRNA transcripts. DNA sequence of IVT templates spanning T7 Phi2.5 promoter to BspQI restriction site are shown. T7 Phi2.5 promoter is underlined. The 5′ UTRs are highlighted in yellow highlight). The canonical eukaryote Kozak sequence is highlighted in black. The start codon and stop codon of the firefly luciferase (Fluc) gene are shown in italic face and highlighted in grey, respectively. The 3’ UTRs are highlight in blue, followed by a 60-nt Poly-A tail. The BspQI restriction site is highlighted in orange.

**Table S3.**
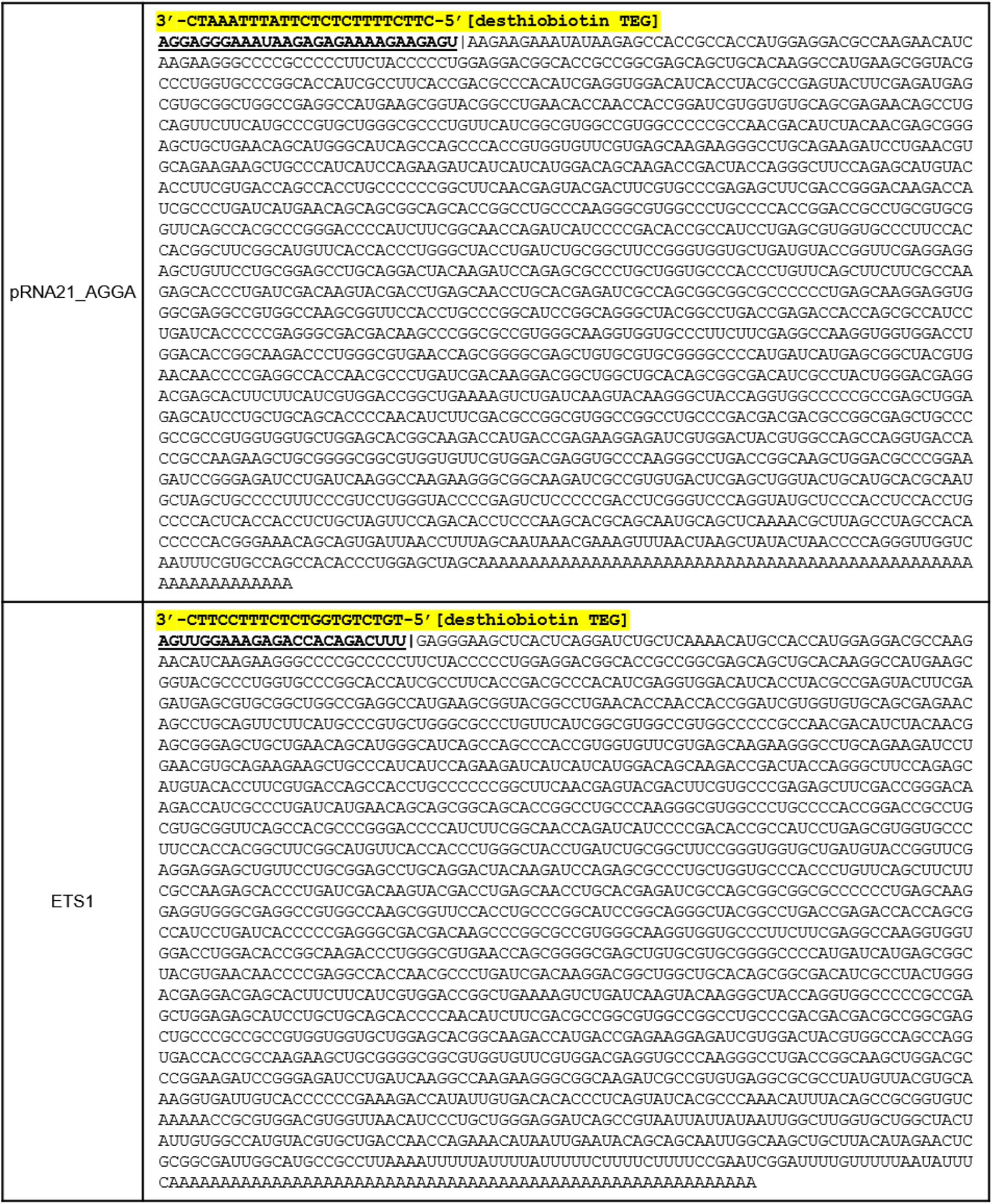
Nucleotide sequence of the model RNA used in this study. The expected nucleotide sequence of the IVT transcripts are shown. The DNA probes used for RNase 4 protection 5′-end analysis are highlighted in yellow. Human RNase 4 hydrolyzes RNA between U and a purine. The expected RNase 4 cleavage site adjacent to the hybridized duplex region is indicated by a “|” sign. The expected 5’ fragments of the IVT transcripts to be analyzed by LC-MS/MS are underlined.

## References

1. Boulias, K. and Greer, E.L. (2022) Biological roles of adenine methylation in RNA. Nature Reviews Genetics 2022 24:3, 24, 143–160.

2. Deng, X., Qing, Y., Horne, D., Huang, H. and Chen, J. (2023) The roles and implications of RNA m6A modification in cancer. Nature Reviews Clinical Oncology 2023 20:8, 20, 507–526.

3. Liu, N., Dai, Q., Zheng, G., He, C., Parisien, M. and Pan, T. (2015) N6-methyladenosine-dependent RNA structural switches regulate RNA–protein interactions. Nature 2015 518:7540, 518, 560–564.

4. Karikó, K., Buckstein, M., Ni, H. and Weissman, D. (2005) Suppression of RNA Recognition by Toll-like Receptors: The Impact of Nucleoside Modification and the Evolutionary Origin of RNA. Immunity, 23, 165–175.

5. Sahin, U., Muik, A., Derhovanessian, E., Vogler, I., Kranz, L.M., Vormehr, M., Baum, A., Pascal, K., Quandt, J., Maurus, D., et al. (2020) COVID-19 vaccine BNT162b1 elicits human antibody and TH1 T cell responses. Nature, 586, 594–599.

6. Adams, J.M. and Cory, S. (1975) Modified nucleosides and bizarre 5ʹ-termini in mouse myeloma mRNA. Nature 1975 255:5503, 255, 28–33.

7. Wei, C.M., Gershowitz, A. and Moss, B. (1975) N6, O2’-dimethyladenosine a novel methylated ribonucleoside next to the 5’ terminal of animal cell and virus mRNAs. Nature, 257, 251–253.

8. Wei, C.M., Gershowitz, A. and Moss, B. (1975) Methylated nucleotides block 5ʹ terminus of HeLa cell messenger RNA. Cell, 4, 379–386.

9. Werner, M., Purta, E., Kaminska, K.H., Cymerman, I.A., Campbell, D.A., Mittra, B., Zamudio, J.R., Sturm, N.R., Jaworski, J. and Bujnicki, J.M. (2011) 2ʹ-O-ribose methylation of cap2 in human: function and evolution in a horizontally mobile family. Nucleic Acids Res., 39, 4756–4768.

10. Despic, V. and Jaffrey, S.R. (2023) mRNA ageing shapes the Cap2 methylome in mammalian mRNA. Nature 2023 614:7947, 614, 358–366.

11. Liu, J.F., Hawley, B.R., Nicholson, L. and Jaffrey, S.R. (2024) Decoding m6Am by simultaneous transcription-start mapping and methylation quantification. Elife, 13.

12. Akichika, S., Hirano, S., Shichino, Y., Suzuki, T., Nishimasu, H., Ishitani, R., Sugita, A., Hirose, Y., Iwasaki, S., Nureki, O., et al. (2019) Cap-specific terminal N 6 -methylation of RNA by an RNA polymerase II– associated methyltransferase. Science (1979)., 363.

13. Boulias, K., Toczydłowska-Socha, D., Hawley, B.R., Liberman, N., Takashima, K., Zaccara, S., Guez, T., Vasseur, J.J., Debart, F., Aravind, L., et al. (2019) Identification of the m6Am Methyltransferase PCIF1 Reveals the Location and Functions of m6Am in the Transcriptome. Mol. Cell, 75, 631–643.e8.

14. Mauer, J., Luo, X., Blanjoie, A., Jiao, X., Grozhik, A. V., Patil, D.P., Linder, B., Pickering, B.F., Vasseur, J.J., Chen, Q., et al. (2017) Reversible methylation of m6 Am in the 5ʹ cap controls mRNA stability. Nature, 541, 371–375.

15. Pandey, R.R., Delfino, E., Homolka, D., Roithova, A., Chen, K.M., Li, L., Franco, G., Vågbø, C.B., Taillebourg, E., Fauvarque, M.O., et al. (2020) The Mammalian Cap-Specific m6Am RNA Methyltransferase PCIF1 Regulates Transcript Levels in Mouse Tissues. Cell Rep., 32.

16. Wang, L., Wu, L., Zhu, Z., Zhang, Q., Li, W., Gonzalez, G.M., Wang, Y. and Rana, T.M. (2023) Role of PCIF1 - mediated 5ʹ-cap N6 -methyladeonsine mRNA methylation in colorectal cancer and anti-PD -1 immunotherapy . EMBO J., 42.

17. Wei, J., Liu, F., Lu, Z., Fei, Q., Ai, Y., He, P.C., Shi, H., Cui, X., Su, R., Klungland, A., et al. (2018) Differential m6A, m6Am, and m1A Demethylation Mediated by FTO in the Cell Nucleus and Cytoplasm. Mol. Cell, 71, 973–985.e5.

18. Shimanski, B., Marin, J.F., Warminski, M., Mckeon, R.M., Kowalska, J., Jemielity, J., Hadden-Perilla, J.A. and Mugridge, J.S. RNA demethylase FTO uses conserved aromatic residues to recognize the mRNA 5ʹ cap and promote efficient m 6 A m demethylation. 10.1101/2025.05.09.653100.

19. Sendinc, E., Valle-Garcia, D., Dhall, A., Chen, H., Henriques, T., Navarrete-Perea, J., Sheng, W., Gygi, S.P., Adelman, K. and Shi, Y. (2019) PCIF1 Catalyzes m6Am mRNA Methylation to Regulate Gene Expression. Mol. Cell, 75, 620–630.e9.

20. Zhuo, W., Sun, M., Wang, K., Zhang, L., Li, K., Yi, D., Li, M., Sun, Q., Ma, X., Liu, W., et al. (2022) m6Am methyltransferase PCIF1 is essential for aggressiveness of gastric cancer cells by inhibiting TM9SF1 mRNA translation. Cell Discovery 2022 8:1, 8, 1–17.

21. Fan, H., Sakuraba, K., Komuro, A., Kato, S., Harada, F. and Hirose, Y. (2003) PCIF1, a novel human WW domain-containing protein, interacts with the phosphorylated RNA polymerase II. Biochem. Biophys. Res. Commun., 301, 378–385.

22. Sun, H., Zhang, M., Li, K., Bai, D. and Yi, C. (2019) Cap-specific, terminal N 6-methylation by a mammalian m6Am methyltransferase. Cell Res., 29, 80–82.

23. Zhuo, W., Sun, M., Wang, K., Zhang, L., Li, K., Yi, D., Li, M., Sun, Q., Ma, X., Liu, W., et al. (2022) m6Am methyltransferase PCIF1 is essential for aggressiveness of gastric cancer cells by inhibiting TM9SF1 mRNA translation. Cell Discov., 8.

24. An, H., Hong, Y., Goh, Y.T., Koh, C.W.Q., Kanwal, S., Zhang, Y., Lu, Z., Yap, P.M.L., Neo, S.P., Wong, C.M., et al. (2024) m6Am sequesters PCF11 to suppress premature termination and drive neuroblastoma differentiation. Mol. Cell, 84, 4142–4157.e14.

25. Gao, S., Zhou, J., Hu, Z., Zhang, S., Wu, Y., Musunuru, P.P., Zhang, T., Yang, L., Luo, X., Bai, J., et al. (2022) Effects of the m6Am methyltransferase PCIF1 on cell proliferation and survival in gliomas. Biochimica et Biophysica Acta (BBA) - Molecular Basis of Disease, 1868, 166498.

26. Wang, L., Wang, S., Wu, L., Li, W., Bray, W., Clark, A.E., Gonzalez, G.M., Wang, Y., Carlin, A.F. and Rana, T.M. (2023) PCIF1-mediated deposition of 5ʹ-cap N6, 2ʹ-O-dimethyladenosine in ACE2 and TMPRSS2 mRNA regulates susceptibility to SARS-CoV-2 infection. Proc. Natl. Acad. Sci. U. S. A., 120, e2210361120.

27. Zhang, Q., Kang, Y., Wang, S., Gonzalez, G.M., Li, W., Hui, H., Wang, Y. and Rana, T.M. (2021) HIV reprograms host m6Am RNA methylome by viral Vpr protein-mediated degradation of PCIF1. Nat. Commun., 12.

28. Tartell, M.A., Boulias, K., Hoffmann, G.B., Bloyet, L.M., Greer, E.L. and Whelan, S.P.J. (2021) Methylation of viral mRNA cap structures by PCIF1 attenuates the antiviral activity of interferon-β. Proc. Natl. Acad. Sci. U. S. A., 118.

29. Catacalos, C., Krohannon, A., Somalraju, S., Meyer, K.D., Janga, S.C. and Chakrabarti, K. (2022) Epitranscriptomics in parasitic protists: Role of RNA chemical modifications in posttranscriptional gene regulation. PLoS Pathog., 18, e1010972.

30. Ben-Haim, M.S., Pinto, Y., Moshitch-Moshkovitz, S., Hershkovitz, V., Kol, N., Diamant-Levi, T., Beeri, M.S., Amariglio, N., Cohen, H.Y. and Rechavi, G. (2021) Dynamic regulation of N6, 2ʹ-O-dimethyladenosine (m6Am) in obesity. Nat. Commun., 12, 1–12.

31. Franco, G., Taillebourg, E., Delfino, E., Homolka, D., Gueguen, N., Brasset, E., Pandey, R.R., Pillai, R.S. and Fauvarque, M.-O. (2023) The catalytic-dead Pcif1 regulates gene expression and fertility in *Drosophila*. RNA, 29, 609–619.

32. Kossykh, V.G., Schlagman, S.L. and Hattman, S. (1993) Conserved sequence motif DPPY in region IV of the phage T4 Dam DNA-[N6-adenine]-methyltransferase is important for S-adenosyl-L-methionine binding. Nucleic Acids Res., 21, 4659–4662.

33. Buratowski, S. (2009) Progression through the RNA polymerase II CTD cycle. Mol. Cell, 36, 541.

34. Doamekpor, S.K., Sanchez, A.M., Schwer, B., Shuman, S. and Lima, C.D. (2014) How an mRNA capping enzyme reads distinct RNA polymerase II and Spt5 CTD phosphorylation codes. Genes Dev., 28, 1323–1336.

35. Chan, S.H., Molé, C.N., Nye, D., Mitchell, L., Dai, N., Buss, J., Kneller, D.W., Whipple, J.M. and Robb, G.B. (2023) Biochemical characterization of mRNA capping enzyme from Faustovirus. RNA, 29, 1803– 1817.

36. Mandell, Z.F., Ujita, A., Henderson, J., Truong, A., Vo, C., Rezvani, F., Abolhassani, N., Lebedev, A., Xu, C., Koukhareva, I., et al. (2025) CleanCap M6 inhibits decapping of exogenously delivered IVT mRNA. Mol. Ther. Nucleic Acids, 36, 102456.

37. Glen Report 35-21: CleanCap® M6.

38. Mirdita, M., Schütze, K., Moriwaki, Y., Heo, L., Ovchinnikov, S. and Steinegger, M. (2022) ColabFold: making protein folding accessible to all. Nature Methods 2022 19:6, 19, 679–682.

39. Jumper, J., Evans, R., Pritzel, A., Green, T., Figurnov, M., Ronneberger, O., Tunyasuvunakool, K., Bates, R., Žídek, A., Potapenko, A., et al. (2021) Highly accurate protein structure prediction with AlphaFold. Nature 2021 596:7873, 596, 583–589.

40. Tuckey, C., Asahara, H., Zhou, Y. and Chong, S. (2014) Protein synthesis using a reconstituted cell-free system. Curr. Protoc. Mol. Biol., 2014, 16.31.1-16.31.22.

41. Wolf, E.J., Dai, N., Chan, S.H. and Corrêa, I.R. (2023) Selective Characterization of mRNA 5ʹ End-Capping by DNA Probe-Directed Enrichment with Site-Specific Endoribonucleases. ACS Pharmacol. Transl. Sci., 6, 1692–1702.

42. Chan, S.H., Tirole, L., Kneller, D., Kelley, T.M., Dai, N. and Robb, G.B. (2023) Co-transcriptional capping using an RNA capping enzyme-T7 RNA polymerase fusion protein. bioRxiv, 10.1101/2023.10.28.564488.

43. Wei, C.M., Gershowitz, A. and Moss, B. (1975) N6, O2’-dimethyladenosine a novel methylated ribonucleoside next to the 5’ terminal of animal cell and virus mRNAs. Nature, 257, 251–253.

44. Akichika, S., Suzuki, T. and Suzuki, T. (2021) Mass spectrometric analysis of mRNA 5ʹ terminal modifications. Methods Enzymol., 658, 407–418.

45. Muthmann, N., Albers, M. and Rentmeister, A. (2023) CAPturAM, a Chemo-Enzymatic Strategy for Selective Enrichment and Detection of Physiological CAPAM-Targets. Angew. Chem. Int. Ed., 62, e202211957.

46. Ishikawa, M., Murai, R., Hagiwara, H., Hoshino, T. and Suyama, K. (2009) Preparation of eukaryotic mRNA having differently methylated adenosine at the 5’-terminus and the effect of the methyl group in translation. Nucleic Acids Symp. Ser. (Oxf)., 10.1093/nass/nrp065.

47. Warminski, M., Trepkowska, E., Smietanski, M., Sikorski, P.J., Baranowski, M.R., Bednarczyk, M., Kedzierska, H., Majewski, B., Mamot, A., Papiernik, D., et al. (2024) Trinucleotide mRNA Cap Analogue N6-Benzylated at the Site of Posttranscriptional m6Am Mark Facilitates mRNA Purification and Confers Superior Translational Properties In Vitro and In Vivo. J. Am. Chem. Soc., 146, 8149–8163.

48. Henderson, J.M., Ujita, A., Hill, E., Yousif-Rosales, S., Smith, C., Ko, N., McReynolds, T., Cabral, C.R., Escamilla-Powers, J.R. and Houston, M.E. (2021) Cap 1 Messenger RNA Synthesis with Co-transcriptional CleanCap® Analog by In Vitro Transcription. Curr. Protoc., 1, e39.

49. Jia, Y. and Patel, S.S. (1997) Kinetic mechanism of GTP binding and RNA synthesis during transcription initiation by bacteriophage T7 RNA polymerase. J. Biol. Chem., 272, 30147–53.

50. Stano, N.M., Levin, M.K. and Patel, S.S. (2002) The +2 NTP binding drives open complex formation in T7 RNA polymerase. J. Biol. Chem., 277, 37292–300.

51. Moroney, S.E. and Piccirilli, J.A. (1991) Abortive products as initiating nucleotides during transcription by T7 RNA polymerase. Biochemistry, 30, 10343–9.

52. Kuzmine, I. and Martin, C.T. (2001) Pre-steady-state kinetics of initiation of transcription by T7 RNA polymerase: a new kinetic model. J. Mol. Biol., 305, 559–66.

53. Ishikawa, M., Murai, R., Hagiwara, H., Hoshino, T. and Suyama, K. (2009) Preparation of eukaryotic mRNA having differently methylated adenosine at the 5’-terminus and the effect of the methyl group in translation. Nucleic Acids Symp. Ser. (Oxf)., 10.1093/nass/nrp065.

54. Roost, C., Lynch, S.R., Batista, P.J., Qu, K., Chang, H.Y. and Kool, E.T. (2015) Structure and thermodynamics of N6-methyladenosine in RNA: A spring-loaded base modification. J. Am. Chem. Soc., 137, 2107– 2115.

55. Roost, C., Lynch, S.R., Batista, P.J., Qu, K., Chang, H.Y. and Kool, E.T. (2015) Structure and thermodynamics of N6-methyladenosine in RNA: A spring-loaded base modification. J. Am. Chem. Soc., 137, 2107– 2115.

56. Spitale, R.C., Flynn, R.A., Zhang, Q.C., Crisalli, P., Lee, B., Jung, J.W., Kuchelmeister, H.Y., Batista, P.J., Torre, E.A., Kool, E.T., et al. (2015) Structural imprints in vivo decode RNA regulatory mechanisms. Nature 2015 519:7544, 519, 486–490.

57. Kuzmine, I., Gottlieb, P.A. and Martin, C.T. (2003) Binding of the Priming Nucleotide in the Initiation of Transcription by T7 RNA Polymerase. Journal of Biological Chemistry, 278, 2819–2823.

58. Roost, C., Lynch, S.R., Batista, P.J., Qu, K., Chang, H.Y. and Kool, E.T. (2015) Structure and Thermodynamics of N6-Methyladenosine in RNA: A Spring-Loaded Base Modification. J. Am. Chem. Soc., 137, 2107–2115.

59. Metkar, M., Pepin, C.S. and Moore, M.J. (2024) Tailor made: the art of therapeutic mRNA design. Nat. Rev. Drug Discov., 23, 67–83.

60. Ohno, H., Akamine, S., Mochizuki, M., Hayashi, K., Akichika, S., Suzuki, T. and Saito, H. (2023) Versatile strategy using vaccinia virus-capping enzyme to synthesize functional 5ʹ cap-modified mRNAs. Nucleic Acids Res., 51, e34.

61. van Dülmen, M., Muthmann, N. and Rentmeister, A. (2021) Chemo-Enzymatic Modification of the 5ʹ Cap Maintains Translation and Increases Immunogenic Properties of mRNA. Angew. Chem. Int. Ed., 60, 13280–13286.

62. Chen, I.M.A., Chu, K., Palaniappan, K., Ratner, A., Huang, J., Huntemann, M., Hajek, P., Ritter, S.J., Webb, C., Wu, D., et al. (2023) The IMG/M data management and analysis system v.7: content updates and new features. Nucleic Acids Res., 51, D723–D732.

63. Mukherjee, S., Stamatis, D., Li, C.T., Ovchinnikova, G., Bertsch, J., Sundaramurthi, J.C., Kandimalla, M., Nicolopoulos, P.A., Favognano, A., Chen, I.M.A., et al. (2023) Twenty-five years of Genomes OnLine Database (GOLD): data updates and new features in v.9. Nucleic Acids Res., 51, D957–D963.

64. The PyMOL Molecular Graphics System, Version 3.0 Schrödinger, LLC.

